# The juvenile-to-adult phase transition in wheat is independent of the winter-spring growth habit regulated by *VRN1*

**DOI:** 10.64898/2026.01.13.699194

**Authors:** Kanata Senoo, Takanori Yoshikawa, Yasir Serag Alnor Gorafi, Shuhei Nasuda

**Affiliations:** Laboratories of Plant Breeding, Graduate School of Agriculture, Kyoto University, Oiwake, Kitashirakawa, Sakyo, Kyoto, 606-8502, Japan; Plant Production Control, Graduate School of Agriculture, Kyoto University, Oiwake, Kitashirakawa, Sakyo, Kyoto, 606-8502, Japan

**Keywords:** juvenile-to-adult phase transition, *miR156*, *miR172*, vegetative phase change, vernalization, *VRN1*, wheat, winter-spring growth habit

## Abstract

In plants, the juvenile-to-adult (JA) phase transition occurs during the vegetative stage with drastic morphological and physiological changes. Common wheat (*Triticum aestivum* L.) has a molecular mechanism regulating the duration of vegetative growth in response to cold accumulation, and its sensitivity varies among varieties (winter-spring growth habit) predominantly due to *VRN1* genotypes. However, the association of the growth habit with the JA phase transition remained unclear. Here, we investigated temporal changes in shoot apex and leaf morphology, and in the expression of the JA phase transition regulators *miR156* and *miR172*, in winter and spring varieties and *VRN1* near-isogenic lines (NILs) under controlled growth conditions, and leaf morphology under field growth conditions. Under controlled conditions, the results indicated that the timing of JA phase transition completion varied among spring varieties without association with *VRN1* genotypes. All NILs underwent the JA phase transition at the same timing, and the expression levels of *miR156* and *miR172* were unrelated to *VRN1* expressions during the vegetative stage. The field evaluation of leaf morphology revealed that the phase transition timing was consistent regardless of the sowing timings. These results suggested that the JA phase transition regulatory pathway and the vernalization regulator *VRN1* are independent in wheat.

**Highlight:** This study shows that the timing of the juvenile-to-adult phase transition in wheat is not affected by the expression of *VRN1,* the master regulator of vernalization.

## Introduction

Higher plants undergo transitions at the proper timing through the embryonic, vegetative, and reproductive stages. They also undergo a transition from the juvenile to the adult phase during the vegetative growth stage, called the juvenile-to-adult phase transition or vegetative phase change (hereafter, JA phase transition). We should note that “juvenile” and “adult” do not refer to “vegetative” and “reproductive”, respectively, but the JA phase transition is defined by drastic changes in leaf morphology, shoot architecture, and physiology during the vegetative stage, as described by Poethig (2013).

The characteristics of the JA phase transition have been well investigated in *Arabidopsis thaliana*. For example, juvenile leaves are smaller and rounder without abaxial trichomes. In contrast, adult leaves are larger and more elongated with abaxial trichomes and higher rates of light-saturated photosynthesis (Telfer *et al*., 1997; Lawrence *et al*., 2021; Poethig and Fouracre, 2024). Juvenile leaves adopt a low-cost and quick-return strategy as they are smaller and structurally simpler but require shorter payback times to obtain the equivalent energy spent on leaf construction than adult leaves, which are high-cost and high-return (Lawrence *et al*., 2022). Abiotic and biotic stress tolerance also alters with age. Juvenile Arabidopsis plants display higher tolerance to submergence stress (Bui *et al*., 2020). A juvenile phase-prolonged transgenic line (*35S::MIR156*) shows a higher recovery rate after salt or drought treatment than the wild type (Col-0) and the juvenile phase-shortened line (*35S::MIM156*) (Cui *et al*., 2014). In terms of biotic stress resistance, adult leaves are more resistant to fungal pathogens and herbivorous insects (Mao *et al*., 2017; Xu *et al*., 2018). These findings indicate that the timing of the JA phase transition and the phenotypes expressed in each phase are crucial survival strategies for plants. Therefore, in the context of agriculture, understanding the regulatory mechanisms of the JA phase transition will provide valuable insights into breeding and cultivation strategies for crops in the era of climate change and global warming, where they are subjected to increasingly severe environmental stress (Rivero *et al*., 2022).

Some key regulatory factors of the JA phase transition have been identified. Among them, *microRNA156* (*miR156*) is a master regulator of the JA phase transition. *miR156* is highly expressed in both leaves and shoot apices during the juvenile phase and is gradually reduced as plant growth and development progress (Wu *et al*., 2009; Xu *et al*., 2016). *miR156* directly targets and inhibits *SQUAMOSA PROMOTER BINDING PROTEIN-LIKE* (*SPL*) genes. Therefore, their expression levels increase throughout development, inducing adult-phase-specific traits (Wu and Poethig, 2006; Wu *et al*., 2009). *SPL*s positively regulate the expression of another microRNA, *miR172*, which targets *APETALA2-LIKE* (*AP2L*) genes, and therefore, the expressions of *miR172* are elevated in the adult phase while those of *AP2Ls* are repressed (Wu *et al*., 2009). *miR172*-*AP2L* module has minor effects on the regulation of adult traits but major effects on the acquisition of reproductive competence (Zhao *et al*., 2023). The *miR156*-*SPL*-*miR172*-*AP2L* pathway is highly conserved in plant species, including major crops such as rice (Xie *et al*., 2006; Zhu *et al*., 2009; Tanaka *et al*., 2011), maize (Lauter *et al*., 2005; Chuck *et al*., 2007), and wheat (Yao *et al*., 2007; Debernardi *et al*., 2022).

Because the characteristics of each phase are species-specific, it is necessary to determine when and how the JA phase transition occurs in the species of interest. Despite extensive studies in Arabidopsis, rice, and maize (Poethig, 1988; Lawson and Poethig, 1995; Asai *et al*., 2002), little is known about wheat, although it is one of the most important cereal crops in the world. Therefore, we previously investigated the JA phase transition in wheat using a Japanese common wheat (*Triticum aestivum* L.) variety Norin 61 as a standard (Senoo *et al*., 2025). We concluded that wheat initiates the JA phase transition from the 1st to the 2nd leaf stage (LS) and reaches the adult phase before the 4th LS. Juvenile leaves are characterized by a smaller ratio of leaf blade length to width, a rounder leaf tip, more adaxial and abaxial trichomes, and higher and lower expressions of *miR156* and *miR172*, respectively. Its transition timing appears to be earlier than in other Poaceae species such as rice, maize, and sorghum (Orkwiszewski and Poethig, 2000; Itoh *et al*., 2005; Hashimoto *et al*., 2019), raising the question of whether regulatory mechanisms specific to wheat exist.

In a normal growth environment in the temperate zone, wheat is exposed to cold temperatures (winter) during early vegetative growth. Therefore, wheat possesses molecular mechanisms to accept cold accumulation as a signal for the initiation of reproductive growth. This trait is known as vernalization. Three *VERNALIZATION* genes, namely *VRN1*, *VRN2*, and *VRN3*, were identified as its key players (Yan *et al*., 2003; Yan *et al*., 2004; Yan *et al*., 2006). *VRN1* is a master regulator of the vernalization and encodes a MADS-box transcription factor, an ortholog of the Arabidopsis floral meristem identifier *APETALA1* (*AP1*) (Mandel and Yanofsky, 1995; Yan *et al*., 2003). *VRN2* encodes a protein that includes a zinc finger and CCT domain, and its counterpart does not exist in Arabidopsis (Yan *et al*., 2004). *VRN3* is also known as *FLOWERING LOCUS T1* (*FT1*), an Arabidopsis ortholog of *FT* (Kardailsky *et al*., 1999; Kobayashi *et al*., 1999; Yan *et al*., 2006). During vernalization, *VRN1* expression is induced in leaves and shoot apical meristems (SAMs), directly repressing *VRN2* and promoting *VRN3* in leaves (Shimada *et al*., 2009; Chen and Dubcovsky, 2012; Deng *et al*., 2015). *VRN2* represses *VRN3* expression under short-day conditions, and this repression is relieved when *VRN1* expression is induced by cold (Hemming *et al*., 2008).

As a result of past breeding efforts, varieties that do not require vernalization, or only require partial vernalization, have been created. The extent of the vernalization requirement is called the winter-spring growth habit. Varieties with a winter growth habit (winter varieties) require vernalization, while those with a spring growth habit (spring varieties) do not. The *VRN1* genotype predominantly determines growth habits. Only one dominant spring-type allele (*Vrn1*), which has insertions, deletions, or single-nucleotide polymorphisms (SNPs) in the promoter region and/or large deletions in the 1st intron compared to the winter-type allele (*vrn1*), is sufficient for the expression of spring growth habit (Fu *et al*., 2005; Zhang *et al*., 2012). This is because the expression of spring-type *Vrn1* is induced early even in the absence of vernalization treatment (Trevaskis *et al*., 2003). Diversity in vernalization requirements has contributed to the success of common wheat cultivation across a wide range of areas worldwide.

Previous studies implied associations between vernalization and the JA phase transition in wheat. Debernardi *et al*. (2022) reported that *VRN1* would positively and negatively regulate *miR172* and *AP2L*, respectively. They also compared *miR156* levels between a spring tetraploid wheat variety Kronos, which has a spring allele on *VRN-A1*, and a mutant line of Kronos (winter Kronos) having a loss-of-function *vrn-A1*. They revealed that the expression patterns both in the 1st leaf of a one-week-old plant and in the 7th leaf of a five-week-old plant were not different between the winter- and spring-type lines. However, JA phase transition would have occurred between these stages, and therefore, the expression analysis at shorter intervals is required to confirm the relationship between *VRN1* and *miR156*. Recently, Liu *et al*. (2026) showed that *VRN1* or its downstream targets upregulate *SPL13* expression, and that *SPL4* can physically interact with the *VRN1* promoter, leading to the coordinated regulation of flowering. Another study in a perennial plant *Cardamine flexuosa* demonstrated that *miR156* and *miR172* levels negatively and positively, respectively, affect the sensitivity to vernalization (Zhou *et al*., 2013). Another report in *Arabis alpina* also supports the idea that high levels of *miR156* prevent the cold-temperature response for flowering (Bergonzi *et al*., 2013). These results suggest potential molecular interactions between the JA phase transition and vernalization pathways. However, these relationships remain unclear.

In this study, we aimed to determine whether the timing of the JA phase transition differs between varieties with winter and spring growth habits and whether this difference can be attributed to *VRN1*-regulated *miR156* or *miR172* expression. To this end, changes in leaf morphology and the expression of *miR156* and *miR172* were investigated in six common wheat varieties, including three winter and three spring varieties each, and *VRN1* near-isogenic lines (NILs) in a controlled environment with warm temperatures and long-day conditions. The results indicated that the timing of the JA phase transition varied among spring varieties. However, *VRN1* would not contribute to the regulation of JA phase transition since *VRN1* expression levels were not associated with the phase transition timing in either the six varieties or the *VRN1* NILs. We attribute the diversity in the JA phase transition among the spring varieties to genetic factors other than *VRN1*. The gene expression analysis revealed that *miR156* and *miR172* regulation are independent of *VRN1* during vegetative growth stage. Additionally, measuring leaf morphological traits under two field conditions, winter-and spring-sowing, showed that the timing of the JA phase transition was comparable between the two conditions, despite differences in *VRN1* expression levels, supporting the independence of the JA phase transition from *VRN1*.

## Materials and Methods

### Plant materials and growth conditions

Three winter common wheat (*Triticum aestivum* L., 2n = 6x = 42, genome constitution = AABBDD) varieties, ‘Norin 10’ (N10), ‘Shunyou’ (SNY), and ‘Jagger’ (JGR), and three spring varieties, ‘Norin 61’ (N61), ‘Chinese Spring’ (CS), and ‘Paragon’ (PRG), were used in this study (Table 1). N10, SNY, N61, and CS seeds were provided by NBRP-Wheat, Japan, (https://shigen.nig.ac.jp/wheat/komugi/) with accession numbers, LPGKU2306, LPGKU2328, LPGKU2305, and LPGKU2320, respectively. JGR (PANG0006) and PRG (PANG0013) seeds are available at the UK Germplasm Resources Unit (SeedStor, https://www.seedstor.ac.uk/).

**Table 1.** List of common wheat varieties used in this study and their genotypes of *VRN1* homoeologs.

| Variety name | Abbreviation | Growth habit | VRN-A1 | VRN-B1 | VRN-D1 |
| --- | --- | --- | --- | --- | --- |
| Norin 10 <sup>a</sup> | N10 | winter | *vrn-A1* | *vrn-B1* | *vrn-D1* |
| Shunyou <sup>a</sup> | SNY | winter | *vrn-A1* | *vrn-B1* | ***Vrn-D1b*** |
| Jagger <sup>b</sup> | JGR | winter | *vrn-A1* | *vrn-B1* | *vrn-D1* |
| Norin 61 <sup>a</sup> | N61 | spring | *vrn-A1* | *vrn-B1* | ***Vrn-D1a*** |
| ChineseSpring <sup>a</sup> | CS | spring | *vrn-A1* | *vrn-B1* | ***Vrn-D1a*** |
| Paragon <sup>b</sup> | PRG | spring | ***Vrn-A1a*** | ***Vrn-B1c*** | *vrn-D1* |

<sup>a</sup> VRN1 genotypes were reported by Mizuno *et al.*, 2022.
<sup>b</sup> VRN1 genotypes were reported by Strejčková *et al.*, 2021.

The genotypes of the *VRN1* homoeologs, *VRN-A1*, *VRN-B1*, and *VRN-D1*, in these varieties are summarized in Table 1 based on previous studies (Strejčková *et al*., 2021; Mizuno *et al*., 2022). Although SNY has a facultative *Vrn-D1b* allele, it has been reported to have a winter growth habit (Mizuno *et al*., 2022). We also confirmed that the winter varieties including SNY had a more delayed heading than the spring varieties in the spring-sowing condition compared to the winter-sowing condition (Supplementary Fig. S1, see the section “Analysis of the JA phase transition in the two growing conditions in the field” for the detailed methods of this measurement). Therefore, SNY was regarded as a winter variety despite its *Vrn-D1* allele. In addition to the six varieties, *VRN1* near-isogenic lines (NILs) with the genetic background of the Japanese common wheat variety Abukumawase were also used. These were developed by Fujita *et al*. (1995) and Seki *et al*. (2007) using Triple Dirk NILs (Pugsley, 1972). They possess one or no spring allele among the *VRN1* homoeologs. Ab(A1), Ab(B1), and Ab(D1) have spring-tyle alleles, namely *Vrn-A1a*, *Vrn-B1a*, and *Vrn-D1a*, respectively, whereas Ab(w) does not have spring alleles. Ab(D4) has a spring allele on the *VRN-D4* locus, which is a functional copy of *VRN-A1* predominantly found in South Asian accessions (Kippes *et al*., 2015).

To promote seed germination, seeds were imbibed in water for one day at 25□, followed by cold exposure at 5□ for two days on several layers of paper towels in a plastic box. Germinated seeds were sown in pots [9.2 cm (l) × 9.2 cm (w) × 12.0 cm (h), volume = 0.52 L] (Nippon Polypot Hanbai, Gifu, Japan) filled with culture soil [red granular soil (“Akadamatsuchi”, Plantation Iwamoto, Ibaraki, Japan): compost (“Sumirin-comparu”, Sumitomo Forestry Landscaping, Osaka, Japan): granular calcium silicate (Shimizu Industrial, Gifu, Japan) = 36:30:1]. Plants were grown at 22.8□ during the day and 20.5□ at night on average under a 16-hour photoperiod with a photosynthetic photon flux density of 76 μmol m^-2^ s^-1^.

### Analysis of morphological traits

In this study, “leaf position” indicates the node number on the main stem where each leaf (1st leaf, 2nd leaf, etc.) is attached. We also used “leaf stage (LS)” to describe the plant’s developmental stage. It refers to the developmental time of plants when each leaf on the main stem is fully developed, marked by the appearance of a lamina joint on the leaf (Senoo *et al*., 2025); for example, it can be described that a plant is at the 2nd LS when its 2nd leaf is fully developed.

For the observation of shoot apices, all leaves were removed using a tweezer and a razor. The exposed shoot apices were observed using a stereomicroscope (Stereomicroscope System SZX7, Olympus, Tokyo, Japan) with a USB digital camera WARYCAM-EL310 (WRAYMER, Osaka, Japan). The color, brightness, and contrast of the photograms were adjusted, and scale bars were added using ImageJ v1.54g (Schneider *et al*., 2012). Three to five biological replicates were used for each variety and NIL.

Leaf size-related traits, including leaf blade length, leaf blade width, and leaf sheath length, were measured using a ruler when each leaf was fully developed. The ratios of leaf blade length to width (length/width) and leaf blade length to leaf sheath length (blade/sheath) were calculated. To facilitate comparison of changes among varieties, measured values from the 1st to the 4th leaves were standardized within each variety by leaf position to have a mean of 0 and a standard deviation of 1 (Z-scores). Five biological replicates were used for each variety, except JGR (n = 4), and six replicates for each NIL.

To measure leaf trichome density, the middle parts of the leaf blades were sampled and fixed in PFA [4% w/v paraformaldehyde and 1% Triton X in PBS(-)]. Four independent fields of view were observed on both the adaxial and abaxial surfaces using a fluorescence microscope BX61 (Olympus, Tokyo, Japan) with 422 nm wavelength light and DAPI filter (Supplementary Fig. S2A), and photographs were taken with a CCD camera DP80 (Olympus, Tokyo, Japan). To reduce manual labor, trichome numbers were first automatically counted using OpenCV v4.12.0 (Bradski, 2000) in Python v3.13.5 (Van Rossum and Drake, 2009), and annotated figures were generated. The source code was written with Microsoft Copilot (https://copilot.microsoft.com/) and is available on GitHub (https://github.com/senookanata/Wheat-leaf-trichome-count). False-positive and false-negative trichomes were manually counted, and the true trichome count was obtained. The automatic counting accuracy was the highest for the PRG samples and the lowest for N61 because of its low trichome density and small trichome size (Supplementary Fig. S2B). Five biological replicates were used for each variety at each leaf position.

Leaf tip angles were measured using the following methodology. The leaf blade shape was scanned using a flatbed scanner DS-50000 (Epson, Nagano, Japan). The leaf tip angle was defined as the angle formed by two straight lines: one connecting the leaf tip and the point on the leaf margin located 1.5 mm below the tip (Fig. 3B). The angles were measured using ImageJ v1.54g. Five biological replicates were used for each variety at each leaf position.

### Quantification of the expression levels of *miR156*, *miR172*, and *VRN1*

The middle parts of the leaf blades (approximately 100 mg) were sampled from the uppermost fully developed leaf blades at each LS. The tissues were quickly frozen in liquid nitrogen and homogenized using a mortar and pestle. Total RNA was extracted using TRIzol Reagent (Thermo Fisher Scientific, Waltham, MA, USA) according to the manufacturer’s protocol, followed by DNase I (Takara, Shiga, Japan) treatment. The first strand cDNA was synthesized using SuperScript III Reverse Transcriptase (Thermo Fisher Scientific) with stem-loop primers for *miR156*, *miR172*, and the inner control *snoR101* as previously reported (Debernardi *et al*., 2022) and with oligo(dT)_20_ primer for *VRN1* and the inner control *GLYCERALDEHYDE-3-PHOSPHATE DEHYDROGENASE* (*GAPDH*) (TraesCS6A03G0586800, TraesCS6B03G0715800, and TraesCS6D03G0485700). 4 μL of 50-times diluted cDNA was used for the real-time qPCR with 1 μL of 10 μM target-specific primer pairs (see Supplementary Table S1) and 5 μL of KOD SYBR qPCR Mix (Toyobo, Osaka, Japan) performed on LightCycler 480 System II (Roche Diagnostics, Rotkreuz, Switzerland). All qPCR primers for miRNAs and *snoR101* were designed by Debernardi *et al*. (2022). *VRN1* primers did not distinguish homoeologous genes, and we quantified their bulked expression (Shimada *et al.,* 2009). *GAPDH* primers were newly developed in this study. The qPCR conditions were as follows: preheating at 98□ for 2 min followed by 45 cycles of denaturation at 98□ for 10 s, annealing at 60□ for 10 s, and extension at 68□ for 30 s. Three biological replicates were used for each line at each stage.

### Analysis of the JA phase transition in the two sowing conditions in the field

The germination-promoting treatment was applied to the seeds as described above. The seeds were sown in 288-cell trays filled with culture soil (H-150, Yanmar, Osaka, Japan) and grown at 22□ with 16 hours of light per day for four or five days. The seedlings were moved with the tray to a nursery house covered with transparent plastic films to acclimate to cold conditions, where it was slightly warmer than the ambient temperature without artificial heating. The seedlings were then transplanted into the field before they reached the 2nd LS. They were planted in a randomized complete block design with three replicates (Supplementary Fig. S3). Each plot consisted of a single row of seven plants from the same accession, spaced 20 cm apart. The distance between adjacent plots was 70 cm. Inorganic fertilizers were applied before transplanting at N = 12, P = 13, and K = 12 g/mm^2^. Organic fertilizers (compost, 375 g/mm^2^) were simultaneously applied, with nutrition provided as N = 9, P = 2, and K = 2 g/mm^2^. The field used in this study was located at Kyoto University, Kyoto, Japan (35.03°N, 135.78°E).

We cultivated the same plant materials at two different timings, namely winter-and spring-sowing conditions, in a single cultivation season, following the above methodology. In the winter-sowing condition, seeds were sown to the cell trays on December 7th, 2024, moved from the growth room with the controlled environments to the nursery house on 12th (9.6□ on average during 12th to 24th), and transplanted on 25th (4.5□ on average during December 25th, 2024 to February 15th, 2025). In the spring-sowing condition, the date of each treatment was 3rd, 7th (11.7□ on average during 7th to 17th), and 18th in March, 2025 (11.4□ on average during March 18th to April 6th). One plant from each replicate (three in total for each accession) was used for serial measurements of leaf morphology (Supplementary Fig. S3). The same individuals were used to record the heading date. Here, the heading was determined by the complete emergence of the first spike.

### Statistical analysis

Statistical tests were performed using the built-in R v4.4.3 package ‘stats’ with the function “t.test(var.equal = equal_var)” for Student’s t-test, “lm()” for linear regression, and “cor.test(method = “Spearman”)” for Spearman’s rank correlation test or the combination of ‘stats’ and the package ‘multcomp v 1.4-28’ with the function “glht(aov())” for Tukey-test (R Core Team, 2025; Hothorn *et al*., 2008). Principal component analysis (PCA) was performed using the stats function “prcomp(scale. = T)” and 95% confidence ellipses of PC1 and PC2 were drawn by ‘ggplot2 v3.5.1’ function “stat_ellipse()” (Wickham, 2016). All graphs were produced using ggplot2 v3.5.1.

## Results

### Morphological changes in the shoot apex and leaf in the six common wheat varieties

We observed shoot apex morphology through the 1st to the 4th LS in the three winter (N10, SNY, and JGR) and three spring (N61, CS, and PRG) common wheat varieties to confirm the duration of the vegetative growth stage. The 1st to the 4th LS of N61, for example, corresponded to 7, 13, 18, and 24 – 25 days after sowing, respectively. The shoot apices of N10, SNY, JGR, and CS consistently showed vegetative forms until the 4th LS, with the expansion of the SAM and the elongation of the tissue beneath it (Fig. 1). N61 showed a vegetative SAM until the 2nd LS, began to transform to a reproductive morphology at the 3rd LS, and completely transitioned to reproductive morphology by the 4th LS. Similarly, the shoot apex of PRG formed reproductive forms from the 3rd LS. These results indicated that N61 and PRG completed the JA phase transition at least before the 4th and 3rd LS, respectively.

**Fig. 1.**
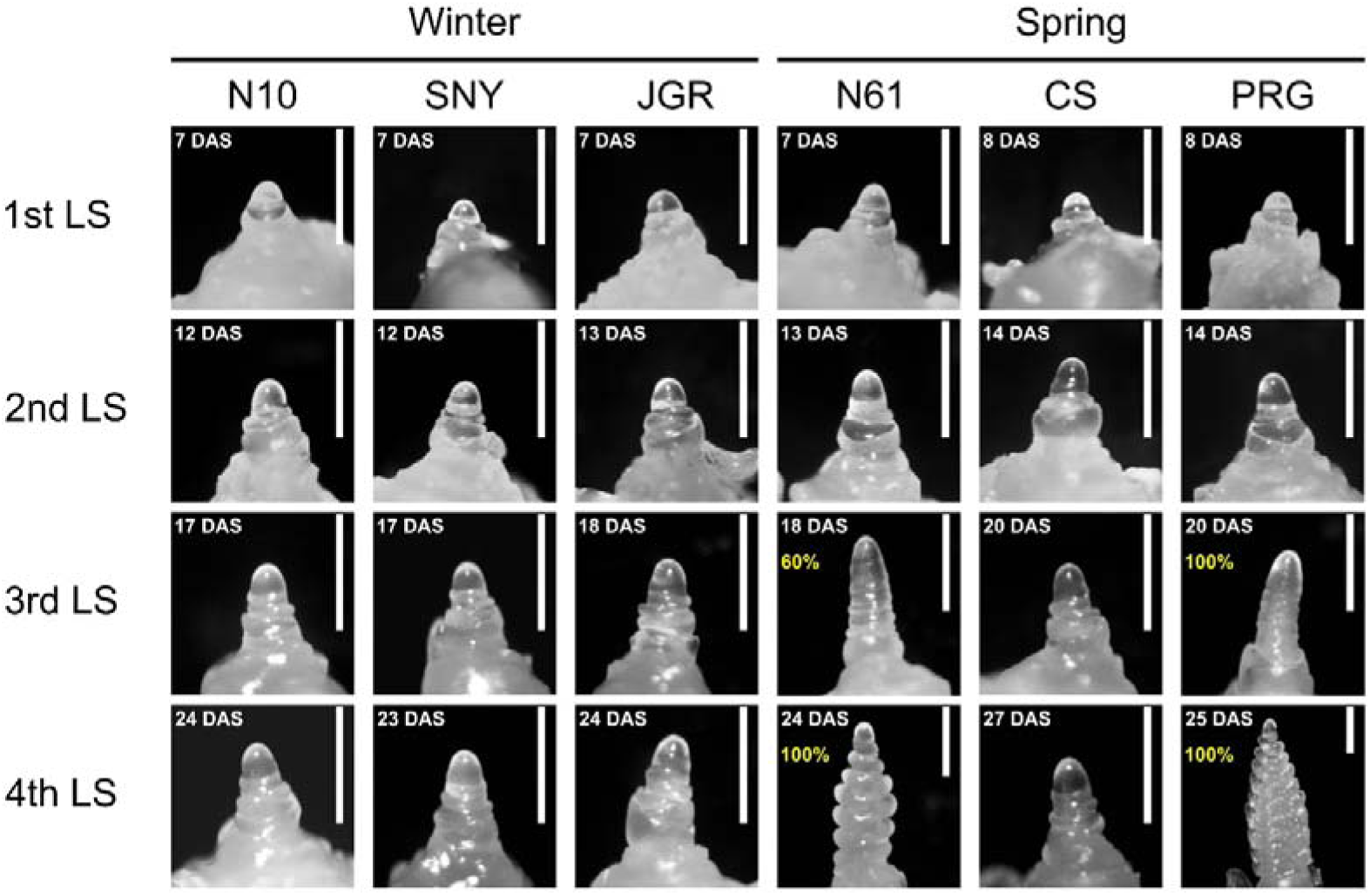
Morphological change in the shoot apices of the six common wheat varieties. “LS” means “leaf stage”. The numbers in the upper left of each picture represent the days after sowing (DAS) of each representative sample. Percentages in figures indicate the proportion of the replicates showing reproductive forms (n = 4, except JGR in the 1st leaf stage (n = 3) and N61, CS, and PRG in the 3rd and 4th leaf stage (n = 5)). The abbreviations of the variety names are as follows: N10 = Norin 10, SNY = Shunyou, JGR = Jagger, N61 = Norin 61, CS = Chinese Spring, and PRG = Paragon. Scale bar = 0.5 mm.

Six leaf size-related traits, namely days to leaf unfoldment, leaf blade length, leaf blade width, ratio of leaf blade length to width (length/width), leaf sheath length, and ratio of leaf blade length to leaf sheath length (blade/sheath) of the 1st to the 4th leaves were measured when each leaf was fully developed (Fig. 2A). Here, we present not only the measured values but also the Z-scores to facilitate the comparison of leaf-position-dependent changes among varieties. The days to the unfolding of each leaf were comparable between the winter and spring varieties. When standardized, differences among the varieties and between the growth habits were diminished, indicating that the phyllochron, the time interval between the emergence of two successive leaves, was stable throughout plant growth in all varieties.

**Fig. 2.**
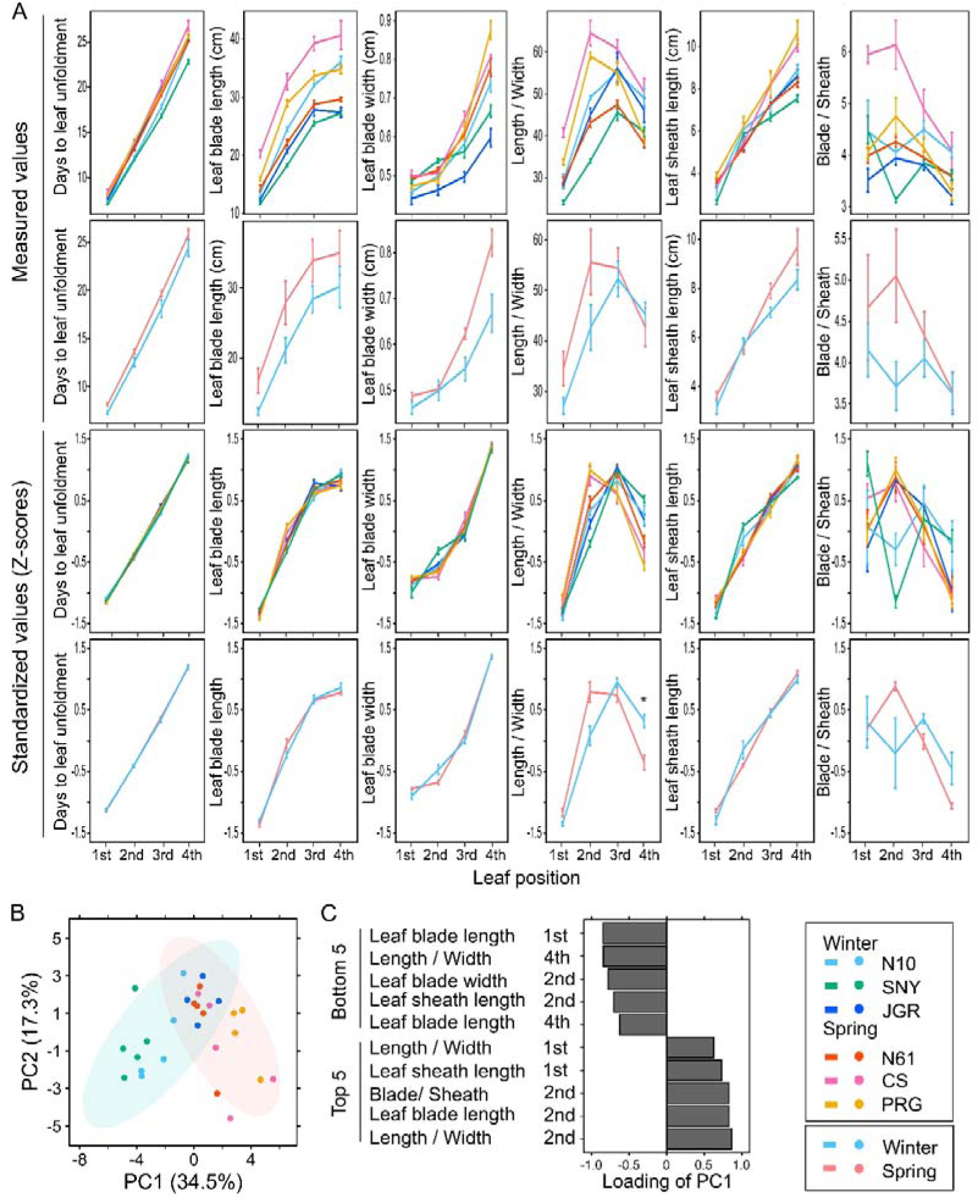
Morphological change in the leaf blades and leaf sheaths of the six common wheat varieties. (A) Measured values (upper two rows) and standardized values (lower two rows) of the six traits related to the leaf size, namely days to leaf unfoldment, leaf blade length, leaf blade width, ratio of leaf blade length to width, leaf sheath length, and ratio of leaf blade length to leaf sheath length, from the 1st to the 4th leaf stage. The top and third figures show the mean and standard error (SE) for each variety. The second and bottom figures show the mean and SE for each growth habit calculated from the mean values of each variety. Asterisk indicates the significant difference between the growth habits by Student’s t-test with Benjamini-Hochberg correction (* p < 0.05). The abbreviations of the variety names are as follows: N10 = Norin 10, SNY = Shunyou, JGR = Jagger, N61 = Norin 61, CS = Chinese Spring, and PRG = Paragon (n = 5 except Jagger (n = 4)). (B) Principal component analysis (PCA) of the Z-scores of the whole traits shown in (A) of all four leaf positions. Each dot represents a biological replicate. The light blue and pink ellipses show the 95% confidence intervals for the winter and spring growth habits, respectively. The proportions of variance explained (PVE) of the PC1 and PC2 are shown in the X- and Y-axis, respectively. (C) The top and bottom five traits with the largest loadings on the PC1.

Leaf blade length was not significantly different between the growth habits, although CS and PRG had longer leaf blades than the others. The Z-scores of the leaf blade length were also insignificant. Spring varieties showed insignificant but large measured values of the leaf blade width of the 4th leaf, while Z-scores were comparable between the growth habits. The unique change in Z-scores was observed in SNY, which was characterized by a larger increase from the 1st to the 2nd leaf than from the 2nd to the 3rd leaf. All varieties showed an increase in the length/width from the 1st to the 2nd leaf, which is an effective indicator of the initiation of the JA phase transition in wheat (Senoo *et al*., 2025). However, CS and PRG showed distinctive changes, reaching their highest values at the 2nd leaf, while the others reached their highest values in the 3rd leaf. In addition, a winter variety, SNY, showed lower values in the 1st to 3rd leaves. Therefore, the spring varieties tended to show larger length/width ratios in the 1st and 2nd leaves, although these differences were not significant. On the other hand, the Z-scores of this ratio showed the similar tendency and were significantly lower in the spring varieties in the 4th leaf. These results reinforced the effectiveness of length/width as a morphological indicator of the timing of the JA phase transition and suggested the possibility of the earlier JA phase transition in spring varieties, particularly in CS and PRG.

No significant differences in the leaf sheath morphology between the growth habits were detected (Fig. 2A). The leaf sheath length of the spring varieties tended to have larger values in the 3rd and 4th leaves, although the difference was not significant. SNY and N10 showed a larger increase in Z-scores from the 1st to the 2nd leaf than in the subsequent leaves. The variation of the blade/sheath among varieties was large in the 1st and the 2nd leaves. CS showed the highest values there, but the difference between growth habits was not significant. Z-scores highlighted a reduction in blade/sheath from the 1st to the 2nd leaf in the two spring varieties, N10 and SNY. Nevertheless, no significant difference was observed in the growth habits at any leaf positions. Overall, changes in leaf size-related traits differed slightly across growth habits. However, the effective indicator, length/width, showed a clear difference in the timing of its maximum, suggesting that spring varieties, particularly CS and PRG, might have undergone an earlier JA phase transition than winter varieties.

To comprehensively observe the differences in leaf morphological changes among varieties, PCA was performed using the Z-scores of all six traits at all leaf positions (Fig. 2B). As a result, PC1, which accounted for 34.5% of the proportion of variance explained (PVE), classified the varieties, where winter and spring varieties showed negative and positive values, respectively. PC1 scores were predominantly influenced by the 1st and 2nd leaves’ traits (Fig. 2C). These results suggested that leaf morphology differed among these varieties according to growth habit.

Since our previous study suggested that changes in leaf trichome density are associated with the JA phase transition (Senoo *et al*., 2025), we also investigated this trait in the middle part of the leaf blades. PRG had the highest density on the 1st to the 3rd leaves on both the adaxial and abaxial surfaces, whereas N61 had the lowest (Fig. 3A). All varieties on the adaxial and abaxial surfaces, except N61, showed an increase from the 1st to the 2nd leaf and a decrease to the 3rd leaf. N61 showed similar changes on the adaxial surface, but almost no trichome was observed on the abaxial surface. There was no significant difference in trichome density between the growth habits on any leaf positions.

**Fig. 3.**
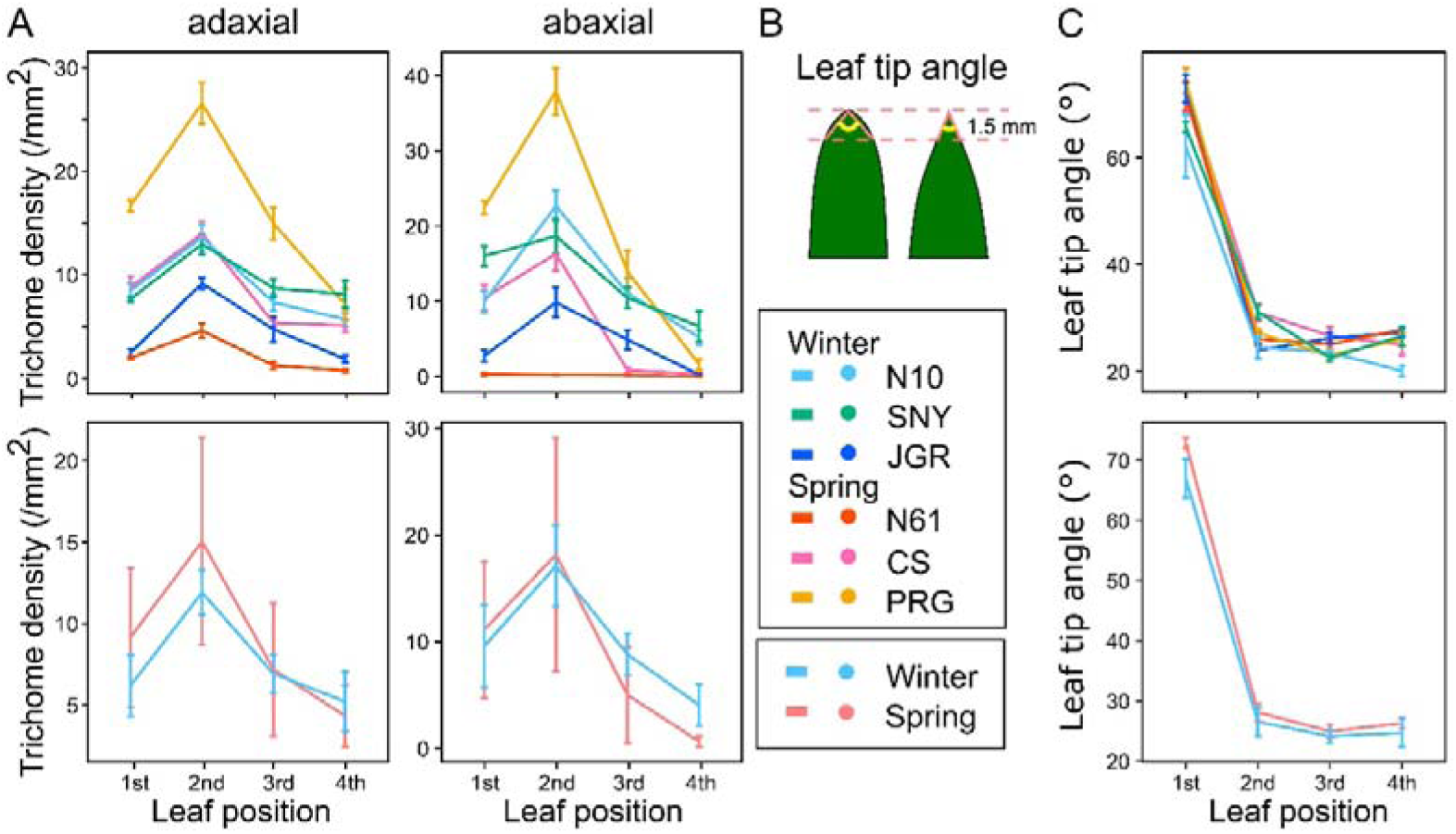
Change in the leaf trichome density and the leaf tip shape of the six common wheat varieties. (A) Trichome density on the adaxial or abaxial side of the middle part of the leaf blades. The top and bottom two figures show the mean and SE for each variety and growth habit, respectively. The abbreviations of the variety names are as follows: N10 = Norin 10, SNY = Shunyou, JGR = Jagger, N61 = Norin 61, CS = Chinese Spring, and PRG = Paragon (n = 5). (B) Definition of leaf tip angle. (C) The results of leaf tip angles shown as mean ± SE for each variety (top) and each growth habit (bottom) (n = 5). No significant difference between the growth habits was detected in either (A) or (C) by Student’s t-test with Benjamini-Hochberg correction.

The leaf tip angle was also measured because a rounder tip was considered a characteristic of juvenile leaves in wheat (Senoo *et al*., 2025). The results revealed that all six varieties had tips that were more than twice as wide at the 1st leaf than the 2nd to the 4th leaves (Fig. 3B and C), suggesting that the 1st leaves of all varieties had juvenile features, whereas the subsequent leaves had transition or adult features. The changes from the 2nd to the 4th leaves were smaller than those between the 1st and the 2nd leaves. Therefore, leaf tip shape could only distinguish between the juvenile and the later phases, but not between the transition and adult phases. Significant differences in growth habits were not observed at any leaf positions. These results suggested that all varieties would begin the JA phase transition from the 1st to the 2nd LS.

In summary, every variety showed large changes in leaf morphology, including length/width, blade/sheath, leaf trichome density, and leaf tip angles, between the 1st and the 2nd leaves (Figs. 2A and 3). Two spring wheat varieties, CS and PRG, reached the peak of the length/width at the 2nd leaf, which was earlier than the other spring variety N61, and all winter varieties (Fig. 2A). These results implied that the completion timing of the JA phase transition tended to be earlier in the two spring varieties, CS and PRG, than in the other varieties.

### Expression changes in the juvenile-to-adult phase transition regulators and their association with *VRN1* in the six varieties

To validate the association between the leaf morphological changes and the expression levels of the JA phase transition regulators, *miR156* and *miR172* levels were quantified by RT-qPCR in uppermost leaf blades through the 1st to the 4th LS. In N10, SNY, JGR, and CS, the *miR156* levels were the highest at the 1st LS and decreased in accordance with the LS, whereas N61 and PRG showed the highest levels at the 2nd LS (Fig. 4A). *miR156* levels were significantly higher at the 1st and 3rd LS in the winter varieties than in the spring varieties. *miR172* levels in all varieties tended to increase as the LS progressed, except JGR showed only minor changes (Fig. 4B). PRG showed a unique pattern, with a large increase in the 2nd LS. However, no significant difference between the growth habits was found at any stages. Taken together with leaf morphology, the higher expression of *miR156* in the winter varieties were highly associated with the later changes of the length/width (Figs. 2A and 4A). In addition, the larger increase in *miR172* level from the 1st to the 2nd LS in PRG was consistent with the earlier completion of the JA phase transition and SAM transition to the reproductive stage (Figs. 1 and 4B).

**Fig. 4.**
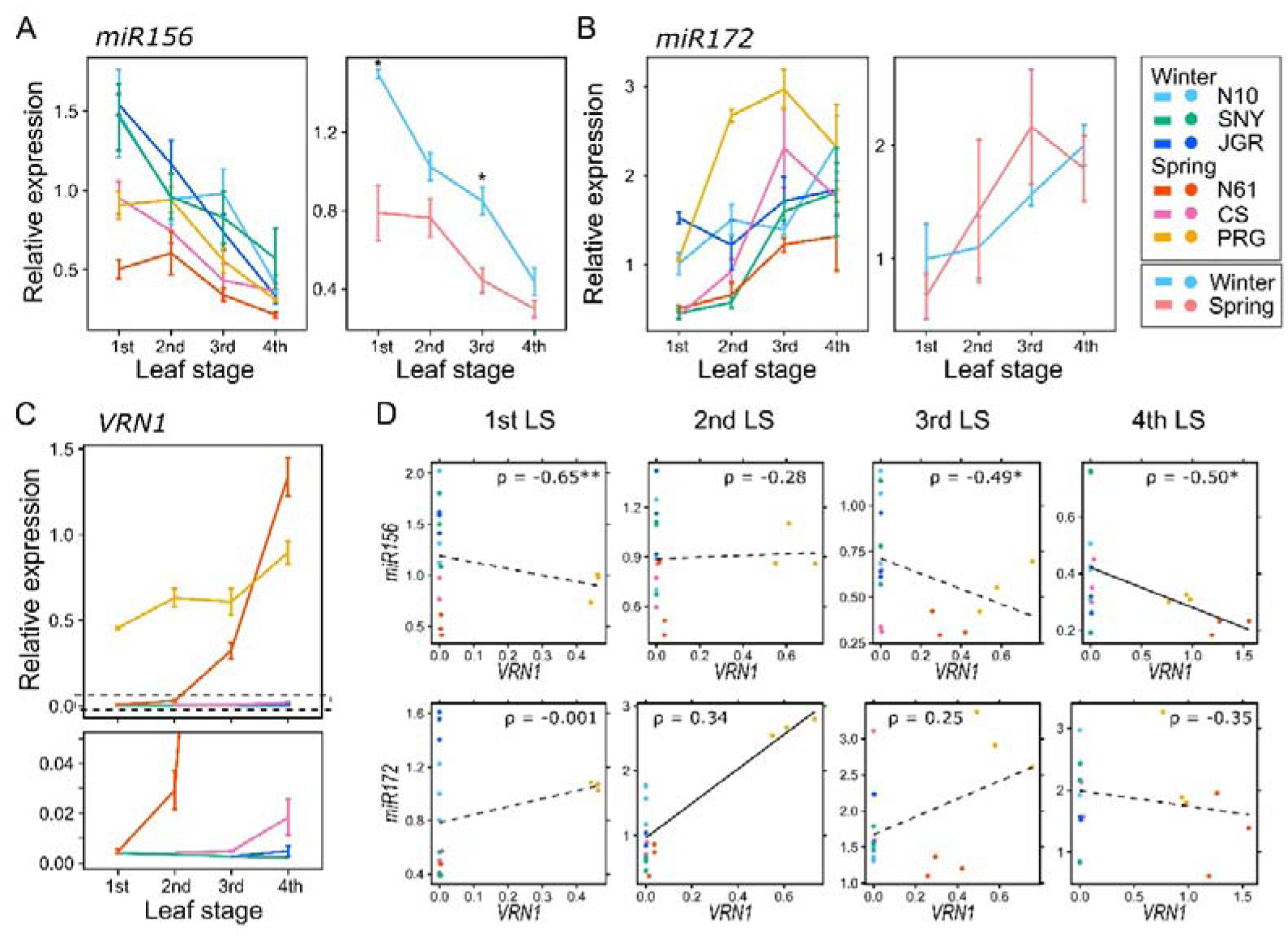
Change in the expression levels of *miR156*, *miR172*, and *VRN1* in the uppermost leaf blade of the six common wheat varieties. (A-C) *miR156* (A) and *miR172* (B) expression levels through the 1st to the 4th leaf stages. The results are shown as mean ± SE for each variety (left) and each growth habit (right). Mean values were compared between winter and spring growth habits using Student’s t-test with Benjamini-Hochberg correction (* indicates p < 0.05). The abbreviations of the variety names are as follows: N10 = Norin 10, SNY = Shunyou, JGR = Jagger, N61 = Norin 61, CS = Chinese Spring, and PRG = Paragon (n =3). (C) *VRN1* expression levels through the 1st to the 4th leaf stages. The results are shown as mean ± SE for each variety. The area enclosed by dotted lines is magnified and presented in the figure below (n = 3). (D) Correlation between *VRN1* and *miR156* (above) or *miR172* (below) expression levels in each leaf stage. Each dot represents a biological replicate. Spearman’s rank correlation coefficients (ρ) are shown above each figure with * (p < 0.05) or ** (p < 0.01). Linear regression is shown in the figure, with solid lines indicating significance at the 5% level and dotted lines indicating non-significance.

To confirm *VRN1*’s involvement, its expression was assessed by RT-qPCR. PRG showed much larger expression even at the 1st LS and kept high levels until the 4th LS (Fig. 4C). N61 showed a large increase from the 2nd to the 3rd and to the 4th LS. On the other hand, the expression was not detected or was quite small until the 4th LS in the three winter varieties and CS. When the expression levels of *miR156* were compared with *VRN1* at each stage, negative Spearman’s rank correlations were observed at the 1st, 3rd, and 4th LS with the statistical significance at the 5% level (Fig. 4D). Comparison of their expression levels across stages showed a significantly negative correlation between *VRN1* and *miR156* at the 3rd and the 4th LS (Supplementary Fig. S4). Regarding the association between *miR172* and *VRN1*, no significant Spearman’s correlation was observed at any stage. However, the 2nd LS showed a significantly positive linear regression (Fig. 4D). This positive regression was driven by PRG’s high *miR172* and *VRN1* levels. Compared across stages, no significant correlation was detected (Supplementary Fig. S5). Although significant correlations between *VRN1* and both *miR156* and *miR172* were observed at several stages, these associations remained unclear because *VRN1* expression levels were too low in many varieties.

Based on the JA phase transition in the six common wheat varieties, it was suggested that they all would initiate the JA phase transition from the 1st to the 2nd LS, but two spring varieties, CS and PRG, would undergo their completion at the 2nd LS earlier than the others at the 3rd LS: they showed the peak in length/width earlier with lower *miR156* expression (Figs. 2A and 4A). However, the effect of *VRN1* genotypes or expression levels on the JA phase transition remained unclear. Therefore, to further confirm their associations, we next attempted to remove noise caused by diverse genetic backgrounds other than *VRN1* using *VRN1* near-isogenic lines (NILs).

### Analysis of the juvenile-to-adult phase transition in *VRN1* near-isogenic lines

Shoot apex morphology was observed from the 1st to the 4th LS in the *VRN1* NILs with a genetic background of a Japanese common wheat variety Abukumawase (See Materials and Methods). The winter line Ab(w) showed vegetative apices until the 4th LS (Fig. 5). The SAMs of the spring lines transitioned to reproductive forms at different stages; Ab(A1) and Ab(D4) were before the 3rd LS and Ab(B1) was around the 3rd to the 4th LS. In Ab(D1), only one out of the four replicates at the 4th LS showed a reproductive apex, suggesting that this line started reproductive growth around the 4th LS. These results indicated that at least Ab(A1) and Ab(D4) reached the adult phase before the 3rd LS. Therefore, changes in traits related to the JA phase transition should be observed during the 1st to the 2nd LS if *VRN1* is involved.

**Fig. 5.**
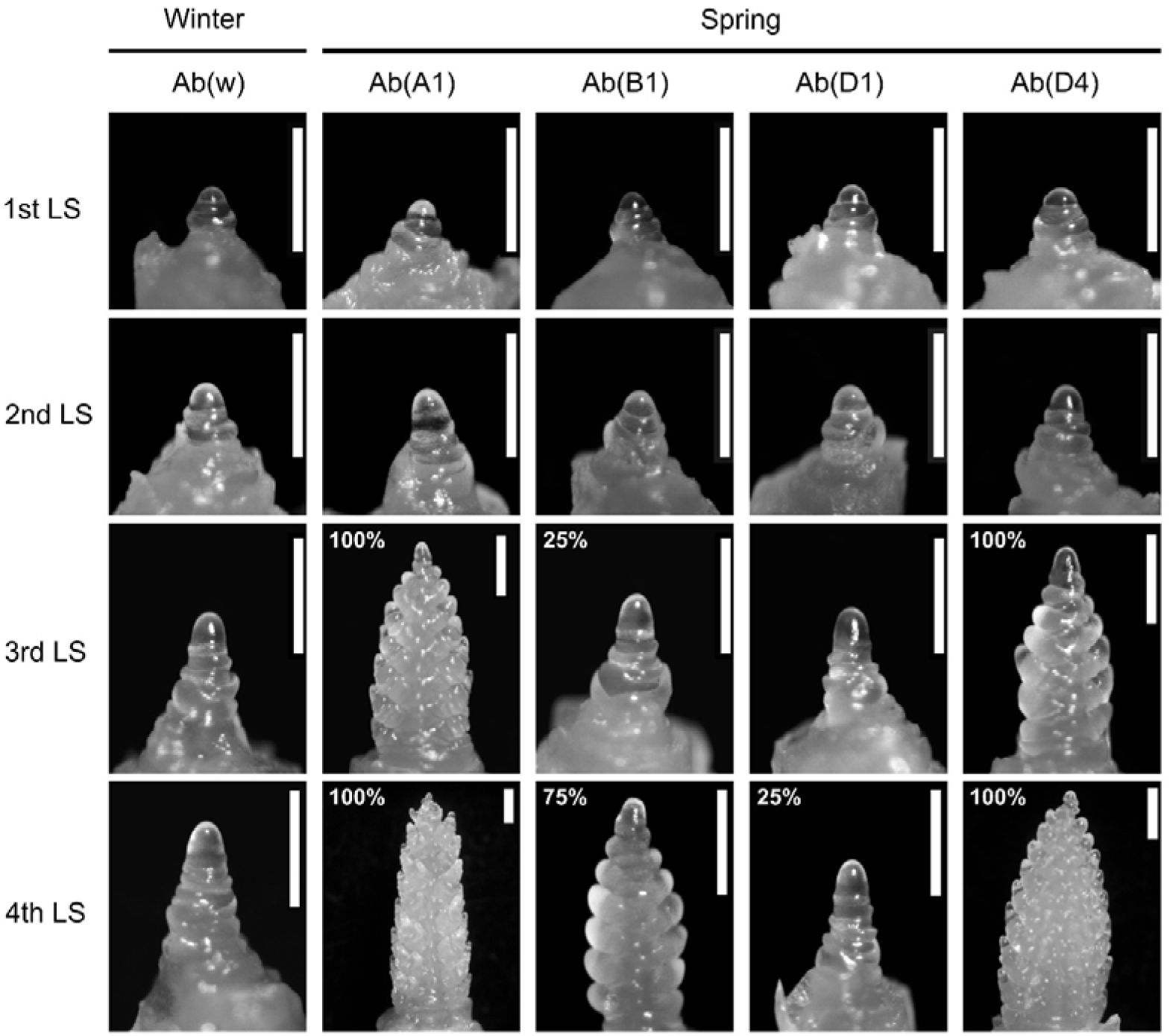
Morphological change in the shoot apices of the *VRN1* near-isogenic lines. “LS” means “leaf stage”. Percentages in figures indicate the proportion of the replicates showing reproductive forms (n = 3 in the 1st leaf stage and n = 4 in the 2nd to the 4th leaf stages). Scale bar = 0.5 mm.

The measurement of leaf size-related traits revealed that the diversity was small among the NILs in the 1st and the 2nd leaves (Fig. 6A). Only Ab(B1), not Ab(A1) or Ab(D4), showed a significant difference in length/width of the 1st leaf. Every line reached the largest values in length/width in the 3rd leaf. Ab(A1) exhibited larger differences from Ab(w) at the 3rd and 4th leaves in days to leaf unfoldment, leaf blade length, length/width, and blade/sheath, when it had already entered the reproductive stage. The small differences in leaf morphology during the vegetative growth stage suggested that there would be no difference in the JA phase transition timing among the *VRN1* NIL.

**Fig. 6.**
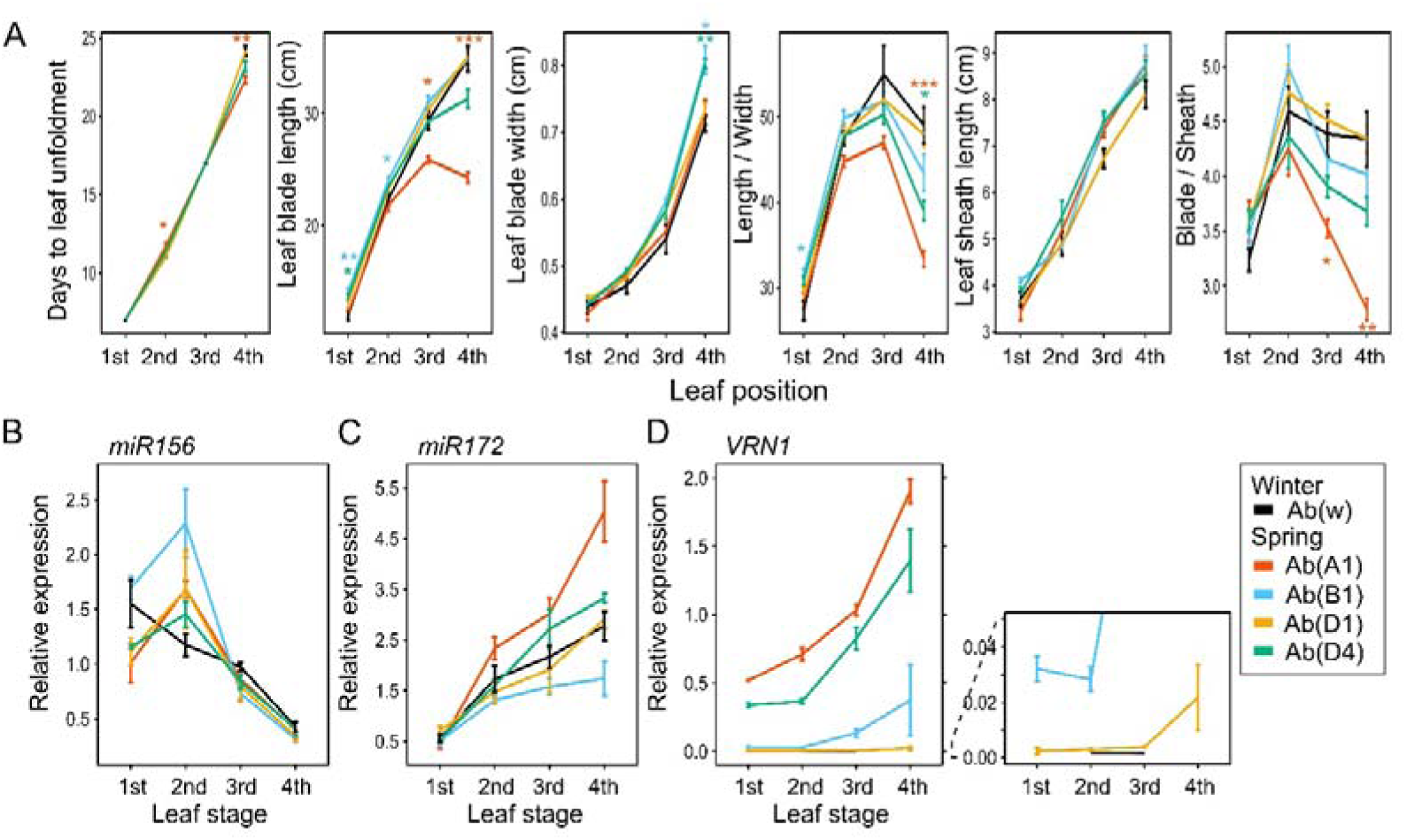
Changes in leaf morphology and expression levels of *miR156*, *miR172*, and *VRN1* in the *VRN1* near-isogenic lines. (A) Measured values of the six traits, days to leaf unfoldment, leaf blade length, leaf blade width, ratio of leaf blade length to width, leaf sheath length, and ratio of leaf blade length to leaf sheath blade, of the 1st to the 4th leaves. Results are shown as the mean and SE for each line (n = 6). (B-D) The expression levels of *miR156* (B), *miR172* (C), and *VRN1* (D) from the 1st to the 4th leaf stages. The results are shown as mean ± SE for each line. (n = 3) A part of the *VRN1* results (D) are magnified and presented in the right figure. Asterisks in (A) show significantly different lines from the winter growth habit line, Ab(w), in each leaf position by Tukey-test (p < 0.05 *, p < 0.01**, and p < 0.001 ***). No significant difference was detected in either (B) or (C) by Tukey-test.

*miR156* and *miR172* expression levels were quantified in the NILs’ leaf blades. The expression of *miR156* was the highest at the 1st LS and gradually decreased in accordance with the LS in Ab(w) (Fig. 6B). On the other hand, its expression increased from the 1st to the 2nd LS and then decreased in the spring lines. *miR172* expressions increased in accordance with the LS in all lines (Fig. 6C). Its levels tended to be higher in Ab(A1) and Ab(D4) at the 3rd and 4th LS. However, no statistically significant differences were detected.

*VRN1* expression levels and their changes varied depending on *VRN1* genotype. Ab(A1) showed a high-level expression even at the 1st LS and a large increase thereafter (Fig. 6D). The expression in Ab(D4) was also high at the 1st LS but slightly lower than Ab(A1). Its level in Ab(B1) was lower than that in the above two lines, and the expression increase was observed after the 2nd LS. Ab(D1) showed lower expression than the other spring lines, and it began to increase after the 3rd LS. In Ab(W), *VRN1* was almost undetected from the 1st to the 4th LS. These results were entirely consistent with the timing of the SAM transition to the inflorescence meristem (Fig. 5).

To clarify the expression relationships between *VRN1* and miRNAs, their levels were compared at each stage or across stages. Only at the 1st LS, *VRN1* showed a significantly negative linear regression against *miR156* (Supplementary Fig. S6). However, *miR156* levels were not associated with *VRN1* levels at any other stages and in any combination of stages (Supplementary Figs. S6 and S7). Therefore, the higher expressions of *miR156* at the 2nd LS in the spring lines would not be attributed to *VRN1*, and its cause remains unclear (Fig. 6B). On the other hand, *miR172* was positively correlated, especially at the 4th LS, where both linear regression and Spearman’s rank correlation showed positive significance (Supplementary Fig. S6). When compared across stages, the positive regressions were observed between *VRN1* at the early stages (1st and 2nd LS) and *miR172* at the later stages (3rd and 4th LS) (Supplementary Fig. S7). The 3rd and 4th LS corresponded to the reproductive stage in Ab(A1) and Ab(D4) (Fig. 5), suggesting that *miR172* regulation by *VRN1* would not have affected the JA phase transition but would have promoted reproductive growth. In conclusion, *VRN1* would contribute to the regulation of *miR172* expression only at the reproductive stage but not to that of *miR156* expression at any stage.

In summary, the timing of the JA phase transition in *VRN1* NILs was not affected by *VRN1* genotypes or expression level. Therefore, the difference in the JA phase transition timing found in the six varieties (Figs. 1 - 4) would be attributed to variations in genetic factors other than *VRN1*.

### JA phase transition in the field conditions and their comparison between the two environmental conditions

Since the completion timing of the JA phase transition varied among the six varieties (N10, SNY, JGR, N61, CS, and PRG) under an artificially controlled environment, we then observed its dynamics in ambient field conditions. We cultivated the same varieties at two independent times, winter and spring, in the same cultivation year (see Materials and Methods). We expected that the different environments, especially temperature and day length, would magnify the difference in the JA phase transition timing among the varieties. In both conditions, only leaf size-related traits were measured.

In the winter-sowing condition, all varieties reached the highest values of length/width in the 2nd leaf, and in the spring-sowing condition, all varieties except JGR also showed the highest in that leaf; JGR showed a slightly higher value in the 1st leaf (Supplementary Fig. S9A). This suggests that all varieties except JGR in the spring condition reached the adult phase at the 2nd LS in both sowing conditions. However, when comparing the length/width values between the sowing conditions, all varieties showed higher values in the 2nd and the 3rd leaves in the winter-sowing condition (Supplementary Fig. S9B). This larger length/width in the winter condition was attributable to the longer leaf blade in the winter-sowing condition, suggesting that the colder temperature or shorter day length might have made the leaf blades longer without affecting the JA phase transition timing.

We also cultivated the *VRN1* NILs in the same conditions. Similar to the six varieties, all NILs in the winter-sowing condition and all NILs except Ab(B1) in the spring-sowing condition showed the highest values of the length/width in the 2nd leaf (Supplementary Fig. S10A). Ab(B1) in the spring-sowing condition showed the reduction of the length/width from the 1st to the 2nd leaf, which was caused by the combination of the smaller increase of the leaf blade length and the larger increase of the leaf blade width than the other NILs. The comparison of the length/width values in each NIL between the sowing conditions revealed that Ab(w), Ab(B1), Ab(D1), and Ab(D4) showed the larger values in the winter-sowing condition until the 3rd leaf, while Ab(A1) showed the larger values in the 1st and the 2nd leaves in the spring-sowing condition (Supplementary Fig. S10B). The reason why Ab(A1) exhibited the opposite trend is still unclear. However, it was supported that *VRN1* expression does not affect the JA phase transition timing because the timings of the length/width peak were same (at the 2nd leaf) among all the NILs in the winter-sowing condition and among most of the NILs in the spring-sowing condition.

## Discussion

This study aimed to elucidate the association between the JA phase transition and the winter-spring growth habit in wheat, and the regulatory mechanisms underlying *VRN1* expression and the phase transition regulators. We compared the JA phase transition-related traits in the winter and spring common wheat varieties and in the *VRN1* NILs. In the analysis of the six varieties including three winter and three spring varieties, all of them showed significant changes in leaf trichome density and leaf tip angle from the 1st to the 2nd leaf (Fig. 3). The ratio of leaf blade length to width was also increased at this timing (Fig. 2A). However, the timing when it hit the highest value was different among varieties; two of the spring varieties, CS and PRG in the 2nd leaf while the other spring variety, N61, and all winter varieties in the 3rd leaf. These results suggested that although all varieties used in this study would have initiated the JA phase transition from the 1st to the 2nd LS, its completion timing would have been earlier in the spring varieties, CS and PRG (2nd LS), than in the other spring variety N61 and all winter varieties, N10, SNY, and JGR (3rd LS). On the other hand, all the *VRN1* NILs analyzed had the highest value at the 3rd leaf, suggesting that *VRN1* genotypes do not affect the JA phase transition (Fig. 6). Taken together, the presence of diversity in the JA completion timing in spring wheat varieties was suggested, which would be attributed to genetic variation(s) other than *VRN1* (Fig. 7A)

**Fig. 7.**
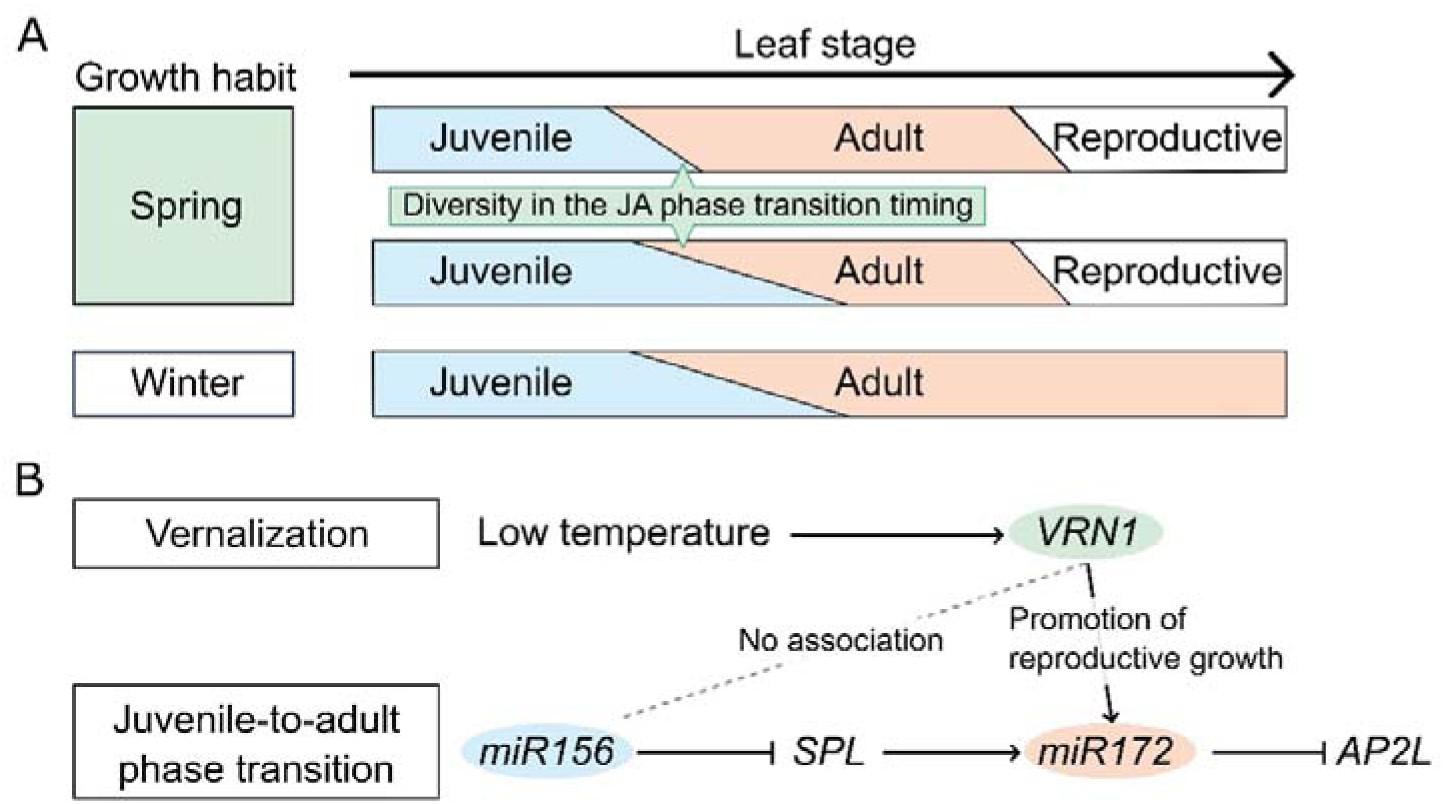
Graphical summary of the relationship between the winter-spring growth habit and the juvenile-to-adult phase transition. (A) Diversity in the juvenile-to-adult phase transition timing found in varieties with spring growth habits. (B) Potential molecular interactions between the vernalization and the juvenile-to-adult phase transition pathways. Solid lines represent the previously reported interactions. The dotted line means that no association exists between *VRN1* and *miR156*. The positive regulation of *miR172* by *VRN1* (solid arrow) previously reported by Debernardi et al. (2022) was also supported in this study, but would only promote reproductive growth, not juvenile-to-adult phase transition.

We did not find any clear correlation between *VRN1* and the JA phase transition regulator *miR156* both among the six varieties and among the *VRN1* NILs (Figs. 4 and 6), indicating that there is no interaction between *VRN1* and *miR156* (Fig. 7B). On the other hand, a positive correlation between *VRN1* and *miR172* was detected among the *VRN1* NILs at the later LSs, which was particularly clear at the 4th LS (Fig. 6 and Supplementary Fig. S6). Previous study also reported that *miR172* is up-regulated by *VRN1*, promoting flowering (Debernardi *et al*., 2022). Another study from the same group demonstrated that the physical interaction of *VRN1* proteins with DELLA proteins, which are negative regulators in the gibberellic acid pathway, can interrupt the repression of SPL3 and SPL4 by DELLA and that *VRN1* also promotes the expression of *SPL13* via a different regulatory mechanism (Liu *et al*., 2026). Considering these findings, the *VRN1*-dependent regulation of *miR172* observed in the present study would have been achieved through VRN1-mediated positive regulation of SPL genes. However, considering that all the NILs showed the same timing of the JA phase transition, it is suggested that the *VRN1*-induced *miR172* would not affect the JA phase transition but would only promote the reproductive growth (Fig. 7B).

Some varieties and all NILs completed the JA transition in the controlled environment at the 3rd LS, while the other varieties were at the 2nd LS. However, under the two field conditions, spring- and winter-sowing, the transition was completed at the 2nd in almost all lines. We attribute this difference between the controlled environment and field conditions, in part, to differences in light intensity, which was about 10 times higher in the field than in the growth room with artificial light. It has been reported that endogenous sucrose and glucose contents promote the JA phase transition through *miR156-SPL9* module (Yang *et al*., 2013; Yu *et al*., 2013; Meng *et al*., 2021). Therefore, the high light intensity in the field might have activated plant photosynthesis, leading to higher sugar accumulation and an earlier transition under field conditions. In addition, the nursing and transplanting of seedlings in the field experiments might have influenced the early-phase transition in the field. In both field conditions, we first grew them in the growth room (22□ with 16 hours of light per day with PPFD of ∼100 μmol m^-2^ s^-1^) until the two or three days before the 1st LS, nursed them in the greenhouse (10 – 12 □ with the natural light) until several days before the 2nd LS, and finally transplanted them to the field. The major differences in leaf morphology between the controlled condition and the field condition were found between the 2nd and the 3rd leaves, where the leaf blade length of all varieties and NILs decreased in the field conditions but increased in the controlled environment (Figs. 2A and 6A; Supplementary Figs. S9A and S10B). Considering these results, the leaf morphology of the 1st and 2nd leaves in field conditions would have been affected mainly by the growth room and greenhouse environments, and that of the 3rd and 4th leaves would have been affected by the ambient field conditions. Environmental factors that change with transplanting, such as decreased temperature and increased nutrient levels, might promote the JA phase transition. To understand the pure effects of the winter- and spring-sowing treatment on the JA phase transition, analysis using direct field sowing rather than nursing will be required.

## Acknowledgement

We thank Dr. Kenji Kato, Okayama University, Japan, for providing the seeds of the NILs. The seeds of the varieties, Norin 10 (LPGKU2306), Shunyou (LPGKU2328), Norin 61 (LPGKU2305), and Chinese Spring (LPGKU2320) were provided by NBRP-Wheat, Japan.

## Supplementary data

The following supplementary data are available at JXB online.

Table S1. Primers used in this study.

Supplementary Fig. S1. Comparison of days to heading between winter- and spring-sowing conditions.

Supplementary Fig. S2. Methodology of the leaf trichome counting.

Supplementary Fig. S3. Experimental design of plant cultivation in the ambient field.

Supplementary Fig. S4. Correlation between *VRN1* and *miR156* expression levels across leaf stages in the six common wheat varieties.

Supplementary Fig. S5. Correlation between *VRN1* and *miR172* expression levels across leaf stages in the six common wheat varieties.

Supplementary Fig. S6. Correlation of the expression levels of *VRN1* and miRNAs in the *VRN1* near-isogenic lines in each leaf stage.

Supplementary Fig. S7. Correlation between *VRN1* and *miR156* expression levels across leaf stages in the *VRN1* near-isogenic lines.

Supplementary Fig. S8. Correlation between *VRN1* and *miR172* expression levels across leaf stage in the *VRN1* near-isogenic lines.

Supplementary Fig. S9. Comparison of leaf morphological changes in the six common wheat varieties between the winter- and spring-sowing conditions in the field.

Supplementary Fig. S10. Comparison of leaf morphological changes in the *VRN1* near-isogenic lines between in the winter- and spring-sowing conditions in the field.

## Author Contribution

Conceptualization: KS, TY, and SN, formal analysis: KS, funding acquisition: SN, investigation: KS, methodology: KS, project administration: SN, resources: KS, supervision: TY and SN, visualization: KS, writing – original draft preparation: KS, and writing – review & editing: TY, YG, and SN.

## Conflict of Interest

No conflict of interest declared.

## Funding

This work was supported in part by JSPS KAKENHI Grant Number JP22K21352 to SN.

## Data Availability

The source code for automatic trichome counting is available on GitHub (https://github.com/senookanata/Wheat-leaf-trichome-count).

## Abbreviations

CS: Chinese Spring
JA: juvenile-to-adult
JGR: Jagger
LS: leaf stage
N10: Norin 10
N61: Norin 61
PRG: Paragon
SAM: shoot apical meristem
SNY: Shunyou
*VRN1*: VERNALIZATION1

## Notes

### Competing Interest Statement

The authors have declared no competing interest.

### Summary of Updates

The description of the last section of the "Results" and the description of the "Discussion" updated. Fig. 7 revised. Supplementary Figs. S1 and S3 newly uploaded.

## References

1. Asai K, Satoh N, Sasaki H, Satoh H, Nagato Y. 2002. A rice heterochronic mutant, *mori1*, is defective in the juvenile-adult phase change. Development 129, 265–273.

2. Bergonzi S, Albani MC, Ver Loren van Themaat E, Nordström KJ, Wang R, Schneeberger K, Moerland PD, Coupland G. 2013. Mechanisms of age-dependent response to winter temperature in perennial flowering of *Arabis alpina*. Science 340, 1094–1097.

3. Bradski G. 2000. The OpenCV Library. Dr. Dobb’s Journal of Software Tools.

4. Bui LT, Shukla V, Giorgi FM, Trivellini A, Perata P, Licausi F, Giuntoli B. 2020. Differential submergence tolerance between juvenile and adult Arabidopsis plants involves the ANAC017 transcription factor. The Plant Jornal 104, 979–994.

5. Chen A, Dubcovsky J. 2012. Wheat TILLING mutants show that the vernalization gene *VRN1* down-regulates the flowering repressor *VRN2* in leaves but is not essential for flowering. PLoS Genetics 8, e1003134.

6. Chuck G, Cigan AM, Saeteurn K, Hake S. 2007. The heterochronic maize mutant *Corngrass1* results from overexpression of a tandem microRNA. Nature Genetics. 39, 544–549.

7. Cui LG, Shan JX, Shi M, Gao JP, Lin HX. 2014. The *miR156-SPL9-DFR* pathway coordinates the relationship between development and abiotic stress tolerance in plants. The Plant Journal 80, 1108–1117.

8. Debernardi JM, Woods DP, Li K, Li C, Dubcovsky J. 2022. MiR172-*APETALA2-like* genes integrate vernalization and plant age to control flowering time in wheat. PLoS Genetics 18, e1010157.

9. Deng W, Casao MC, Wang P, Sato K, Hayes PM, Finnegan EJ, Trevaskis B. 2015. Direct links between the vernalization response and other key traits of cereal crops. Nature Communications. 6, 5882.

10. Fu D, Szucs P, Yan L, Helguera M, Skinner JS, von Zitzewitz J, Hayes PM, Dubcovsky J. 2005. Large deletions within the first intron in *VRN-1* are associated with spring growth habit in barley and wheat. Molecular Genetics and Genomics 273, 54–65.

11. Fujita M, Taniguchi Y, Ujihara K, Sasaki A. 1995. Ear Primordia Development and Stem Elongation of Near-isogenic Lines for Vernalization Requirement in Extremely-Early Maturing Wheat Cultivars (T*riticum aestivum* L.). Vol. 45. Breeding Science, 97–104.

12. Hashimoto S, Tezuka T, Yokoi S. 2019. Morphological changes during juvenile-to-adult phase transition in sorghum. Planta 250, 1557–1566.

13. Hemming MN, Peacock WJ, Dennis ES, Trevaskis B. 2008. Low-temperature and daylength cues are integrated to regulate *FLOWERING LOCUS T* in barley. Plant Physiology 147, 355–366.

14. Hothorn T, Bretz F, Westfall P. 2008. Simultaneous inference in general parametric models. Biometrical Journal 50, 346–363.

15. Itoh J, Nonomura K, Ikeda K, Yamaki S, Inukai Y, Yamagishi H, Kitano H, Nagato Y. 2005. Rice plant development: from zygote to spikelet. Plant and Cell Physiology 46, 23–47.

16. Kardailsky I, Shukla VK, Ahn JH, Dagenais N, Christensen SK, Nguyen JT, Chory J, Harrison MJ, Weigel D. 1999. Activation tagging of the floral inducer *FT*. Science 286, 1962–1965.

17. Kippes N, Debernardi JM, Vasquez-Gross HA, Akpinar BA, Budak H, Kato K, Chao S, Akhunov E, Dubcovsky J. 2015. Identification of the *VERNALIZATION 4* gene reveals the origin of spring growth habit in ancient wheats from South Asia. Proceedings of National Academy of Science of the United States of America 112, E5401–5410.

18. Kobayashi Y, Kaya H, Goto K, Iwabuchi M, Araki T. 1999. A pair of related genes with antagonistic roles in mediating flowering signals. Science 286, 1960–1962.

19. Lauter N, Kampani A, Carlson S, Goebel M, Moose SP. 2005. *microRNA172* down-regulates *glossy15* to promote vegetative phase change in maize. Proceedings of National Academy of Science of the United States of America 102, 9412–9417.

20. Lawrence EH, Springer CJ, Helliker BR, Poethig RS. 2021. MicroRNA156-mediated changes in leaf composition lead to altered photosynthetic traits during vegetative phase change. New Phytologist 231, 1008–1022.

21. Lawrence EH, Springer CJ, Helliker BR, Poethig RS. 2022. The carbon economics of vegetative phase change. Plant, Cell & Environ 45, 1286–1297.

22. Lawson EJR, Poethig RS. 1995. Shoot development in plants: time for a change. Trends in Genetics. 11, 263–268.

23. Liu Q, Zhang L, Zhou Z, Zhang C, Li C, Debernardi JM, Dubcovsky J. 2026. MicroRNA156 and its targeted *SPL* genes interact with the photoperiod, vernalization, and gibberellin pathways to regulate wheat heading time. The Plant Journal 125, e70656.

24. Mandel MA, Yanofsky MF. 1995. A gene triggering flower formation in Arabidopsis. Nature 377, 522–524.

25. Mao YB, Liu YQ, Chen DY, Chen FY, Fang X, Hong GJ, Wang LJ, Wang JW, Chen XY. 2017. Jasmonate response decay and defense metabolite accumulation contributes to age-regulated dynamics of plant insect resistance. Nature Communications 8, 13925.

26. Meng LS, Bao QX, Mu XR, Tong C, Cao XY, Huang JJ, Xue LN, Liu CY, Fei Y, Loake GJ. 2021. Glucose- and sucrose-signaling modules regulate the Arabidopsis juvenile-to-adult phase transition. Cell Reports 36, 109348.

27. Mizuno N, Matsunaka H, Yanaka M, Nakata M, Nakamura K, Nakamaru A, Kiribuchi-Otobe C, Ishikawa G, Chono M, Hatta K, Fujita M, Kobayashi F. 2022. Allelic variations of *Vrn-1* and *Ppd-1* genes in Japanese wheat varieties reveal the genotype-environment interaction for heading time. Breeding Science 72, 343–354.

28. Orkwiszewski JA, Poethig RS. 2000. Phase identity of the maize leaf is determined after leaf initiation. Proceedings of National Academy of Science of the United States of America 97, 10631–10636.

29. Poethig RS, Fouracre J. 2024. Temporal regulation of vegetative phase change in plants. Developmental Cell 59, 4–19.

30. Poethig RS. 1988. Heterochronic mutations affecting shoot development in maize. Genetics 119, 959–973.

31. Poethig RS. 2013. Vegetative phase change and shoot maturation in plants. Current Topics in Developmental Biology 105, 125–152.

32. Pugsley AT. 1972. Additional genes inhibiting winter habit in wheat. Vol. 21. Euphytica, 547–552.

33. R Core Team. 2025. R: A Language and Environment for Statistical Computing. R Foundation for Statistical Computing, Vienna, Austria.

34. Rivero RM, Mittler R, Blumwald E, Zandalinas SI. 2022. Developing climate-resilient crops: improving plant tolerance to stress combination. The Plant Journal 109, 373–389.

35. Schneider CA, Rasband WS, Eliceiri KW. 2012. NIH Image to ImageJ: 25 years of image analysis. Nature Methods 9, 671–675.

36. Seki M, Oda S, Matsunaka H, Hatta, Kouichi, Fujita, Masaya, Hatano T, Otobe-Kiribuchi C, Kawada N, Kato K. 2007. Growth and Yield of Near-Isogenic Wheat Lines Carrying Different Vernalization Response Genes. Vol. 9. Breeding Research: J-STAGE, 125–133.

37. Senoo K, Yoshioka S, Yamamori K, Nasuda S, Yoshikawa T. 2025. Juvenile-to-adult phase transition in a common wheat cultivar Norin 61, and accompanying changes in leaf transcriptome. Plant and Cell Physiology 66, 900–911.

38. Shimada S, Ogawa T, Kitagawa S, Suzuki T, Ikari C, Shitsukawa N, Abe T, Kawahigashi H, Kikuchi R, Handa H, Murai K. 2009. A genetic network of flowering-time genes in wheat leaves, in which an *APETALA1*/*FRUITFULL*-like gene, *VRN1*, is upstream of *FLOWERING LOCUS T*. The Plant Journal 58, 668–681.

39. Strejčková B, Milec Z, Holušová K, Cápal P, Vojtková T, Čegan R, Šafář J. 2021. In-depth sequence analysis of bread wheat *VRN1* genes. International Journal of Molecular Sciences 22.

40. Tanaka N, Itoh H, Sentoku N, Kojima M, Sakakibara H, Izawa T, Itoh J, Nagato Y. 2011. The *COP1* ortholog PPS regulates the juvenile-adult and vegetative-reproductive phase changes in rice. The Plant Cell 23, 2143–2154.

41. Telfer A, Bollman KM, Poethig RS. 1997. Phase change and the regulation of trichome distribution in Arabidopsis thaliana. Development 124, 645–654.

42. Trevaskis B, Bagnall DJ, Ellis MH, Peacock WJ, Dennis ES. 2003. MADS box genes control vernalization-induced flowering in cereals. Proceedings of National Academy of Science of the United States of America 100, 13099–13104.

43. Van Rossum G, Drake FL. 2009. Python 3 Reference Manual. Vol. Scotts Valley, CA. CreateSpace.

44. Wickham H. 2016. ggplot2: Elegant Graphics for Data Analysis. Springer-Verlag New York.

45. Wu G, Park MY, Conway SR, Wang JW, Weigel D, Poethig RS. 2009. The sequential action of miR156 and miR172 regulates developmental timing in *Arabidopsis*. Cell 138, 750–759.

46. Wu G, Poethig RS. 2006. Temporal regulation of shoot development in *Arabidopsis thaliana* by *miR156* and its target *SPL3*. Development 133, 3539–3547.

47. Xie K, Wu C, Xiong L. 2006. Genomic organization, differential expression, and interaction of SQUAMOSA promoter-binding-like transcription factors and microRNA156 in rice. Plant Physiology 142, 280–293.

48. Xu M, Hu T, Zhao J, Park MY, Earley KW, Wu G, Yang L, Poethig RS. 2016. Developmental Functions of miR156-Regulated *SQUAMOSA PROMOTER BINDING PROTEIN-LIKE (SPL)* Genes in *Arabidopsis thaliana*. PLoS Genetics 12, e1006263.

49. Xu YP, Lv LH, Xu YJ, Yang J, Cao JY, Cai XZ. 2018. Leaf stage-associated resistance is correlated with phytohormones in a pathosystem-dependent manner. Journal of Integrative Plant Biology 60, 703–722.

50. Yan L, Fu D, Li C, Blechl A, Tranquilli G, Bonafede M, Sanchez A, Valarik M, Yasuda S, Dubcovsky J. 2006. The wheat and barley vernalization gene *VRN3* is an orthologue of *FT*. Proceedings of National Academy of Science of the United States of America 103, 19581–19586.

51. Yan L, Loukoianov A, Blechl A, Tranquilli G, Ramakrishna W, SanMiguel P, Bennetzen JL, Echenique V, Dubcovsky J. 2004. The wheat *VRN2* gene is a flowering repressor down-regulated by vernalization. Science 303, 1640–1644.

52. Yan L, Loukoianov A, Tranquilli G, Helguera M, Fahima T, Dubcovsky J. 2003. Positional cloning of the wheat vernalization gene *VRN1*. Proceedings of National Academy of Science of the United States of America 100, 6263–6268.

53. Yang L, Xu M, Koo Y, He J, Poethig RS. 2013. Sugar promotes vegetative phase change in *Arabidopsis thaliana* by repressing the expression of *MIR156A* and *MIR156C*. Elife 2, e00260.

54. Yao Y, Guo G, Ni Z, Sunkar R, Du J, Zhu JK, Sun Q. 2007. Cloning and characterization of microRNAs from wheat (*Triticum aestivum* L.). Genome Biology. 8, R96.

55. Yu S, Cao L, Zhou CM, Zhang TQ, Lian H, Sun Y, Wu J, Huang J, Wang G, Wang JW. 2013. Sugar is an endogenous cue for juvenile-to-adult phase transition in plants. Elife 2, e00269.

56. Zhang J, Wang Y, Wu S, Yang J, Liu H, Zhou Y. 2012. A single nucleotide polymorphism at the *Vrn-D1* promoter region in common wheat is associated with vernalization response. Theoretical and Applied Genetics 125, 1697–1704.

57. Zhao J, Doody E, Poethig RS. 2023. Reproductive competence is regulated independently of vegetative phase change in *Arabidopsis thaliana*. Current Biology 33, 487–497.e482.

58. Zhou CM, Zhang TQ, Wang X, Yu S, Lian H, Tang H, Feng ZY, Zozomova-Lihová J, Wang JW. 2013. Molecular basis of age-dependent vernalization in Cardamine flexuosa. Science 340, 1097–1100.

59. Zhu QH, Upadhyaya NM, Gubler F, Helliwell CA. 2009. Over-expression of miR172 causes loss of spikelet determinacy and floral organ abnormalities in rice (*Oryza sativa*). BMC Plant Biology 9, 149.

