## Supplementary File for "The juvenile-to-adult phase transition in wheat is independent of the winter-spring growth habit regulated by *VRN1*"

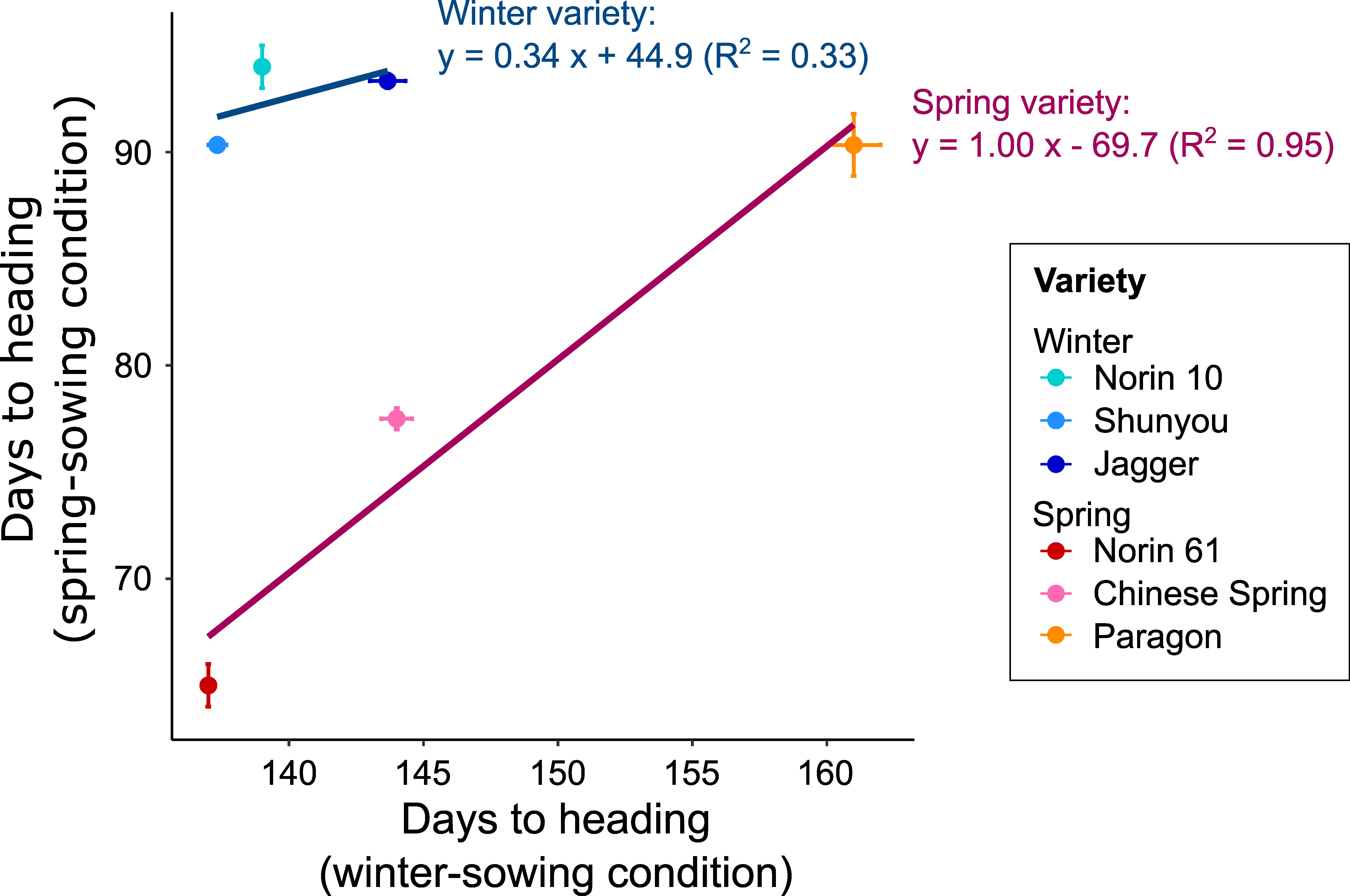

**Supplementary Fig. S1. Comparison of days to heading between winter- and spring-sowing conditions.**

“Days to heading” was defined here as the number of days from the sowing of seed to the complete emergence of a first spike in each plant. Results were shown as mean ± SE (n = 3 in each growth condition except Chinese Spring, which had only two replicates in the spring-sowing condition). Mean values in each variety were used for the linear regression analysis.

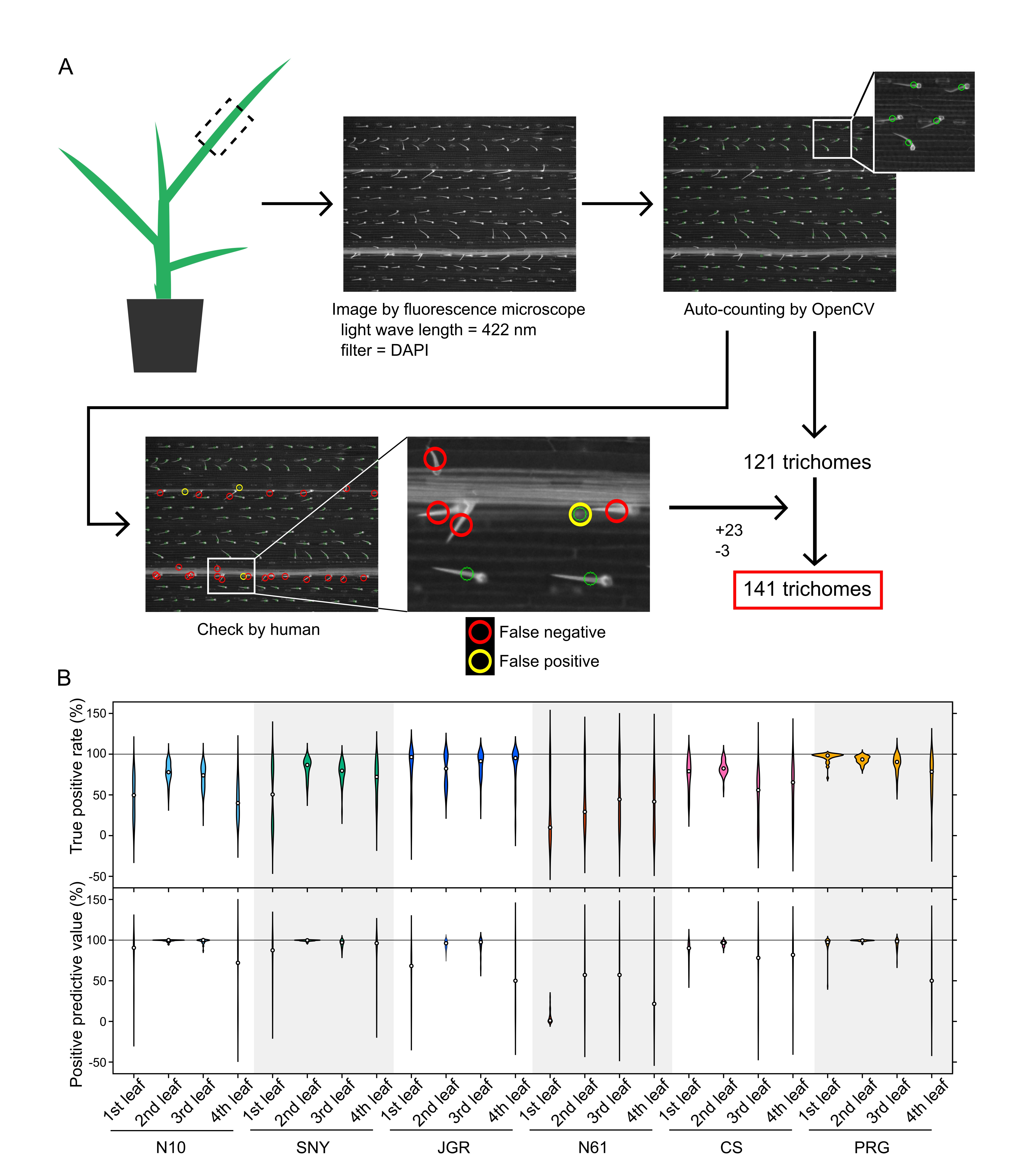

**Supplementary Fig. S2. Methodology of the leaf trichome counting.**

(A) Scheme of the semi-automatic trichome counting. (B) Counting accuracy of the semi-automatic approach. Each violine plot includes 40 data points derived from four microscopic images on two leaf sides (adaxial and abaxial) of five biological replicates. Dots represent medians.

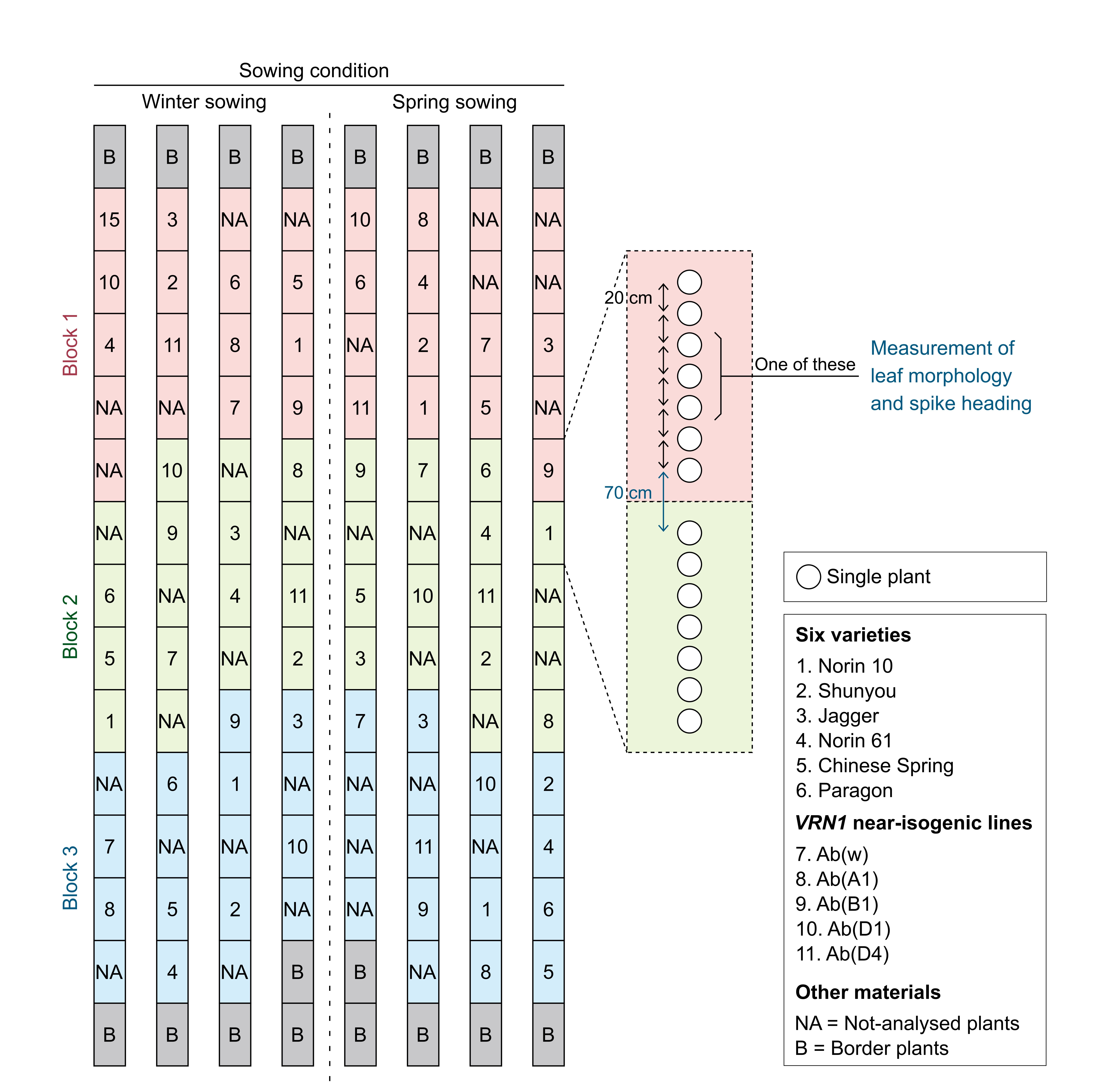

**Supplementary Fig. S3. Experimental design of plant cultivation in the ambient field.**

In each block (Block 1, Block 2, and Block 3), each plot consisting of a single germplasm was randomly arranged. In each plot, seven plants were transplanted with 20 cm intervals. The distance between adjacent plots was 70 cm. One individual from each plot was used for the non-destructive measurements of leaf morphology. Three individuals were used for the destructive sampling of leaves for the gene expression analysis.

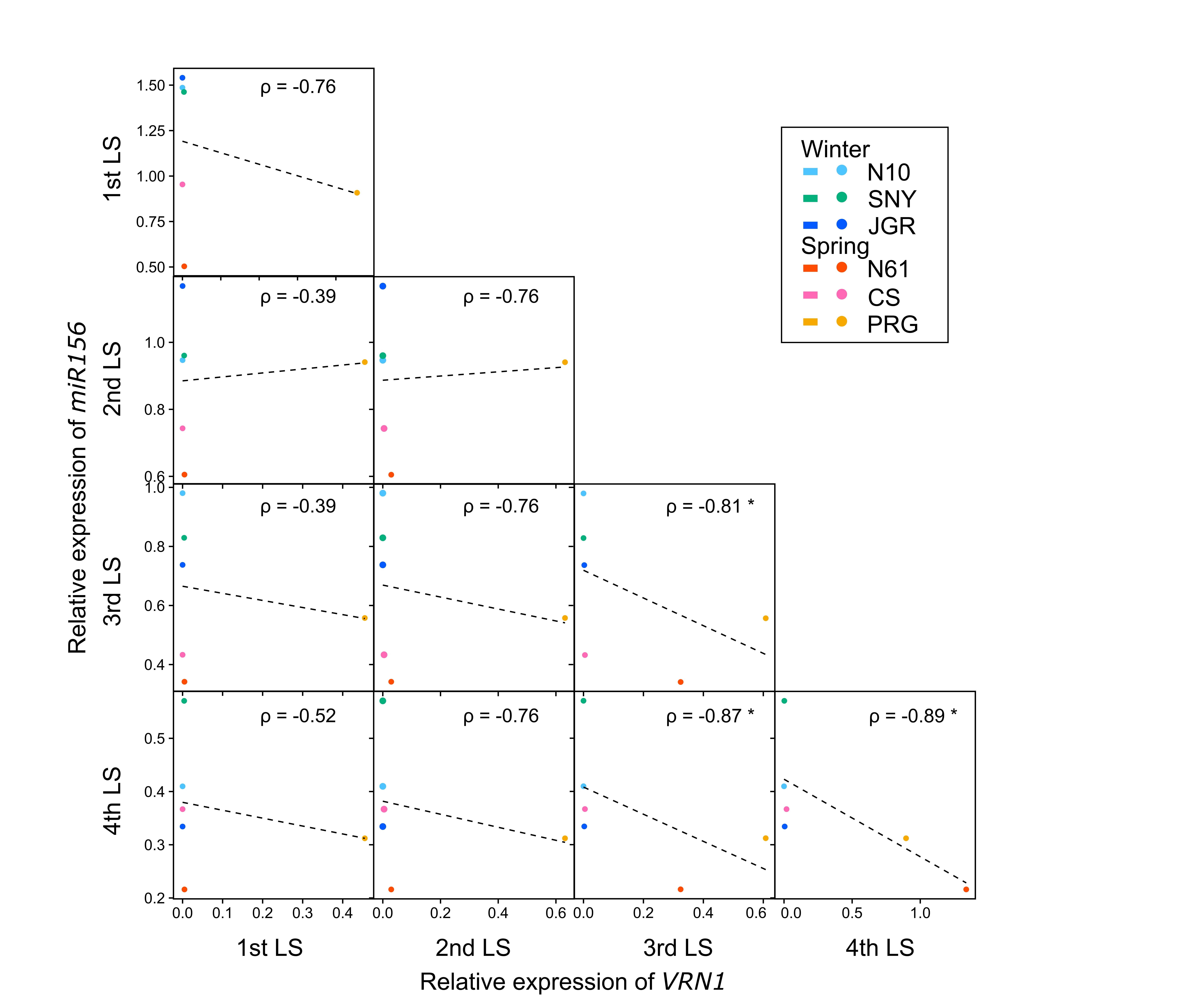

**Supplementary Fig. S4. Correlation between *VRN1* and *miR156* expression levels across leaf stages in the six common wheat varieties.**

Each dot represents the mean value in each variety. Spearman’s rank correlation coefficients (ρ) are shown in each figure with * (p < 0.05). Linear regression is shown in the figure, all of which were not significant at the 5% level.

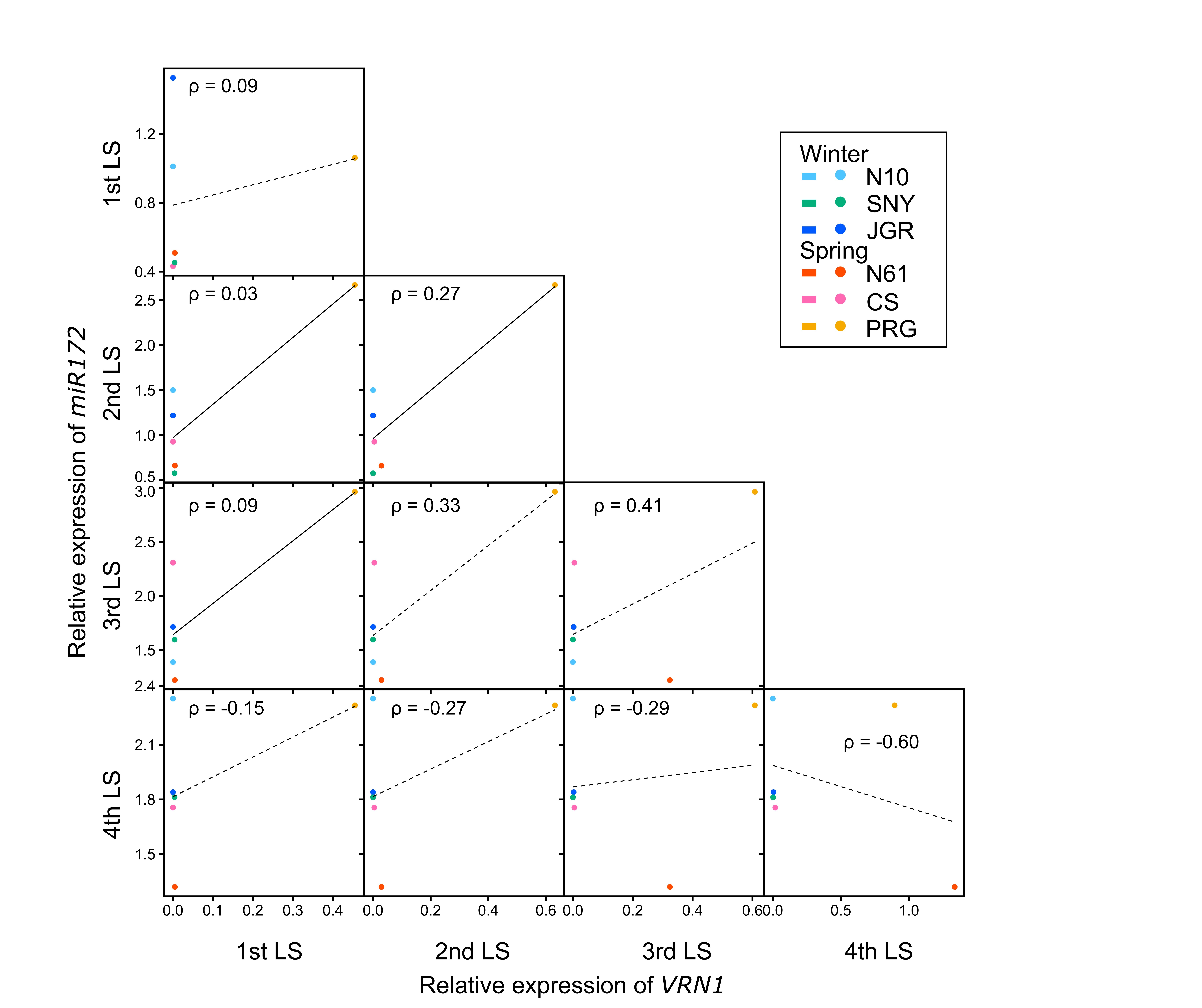

**Supplementary Fig. S5. Correlation between *VRN1* and *miR172* expression levels across leaf stages in the six common wheat varieties.**

Each dot represents the mean value in each variety. Spearman’s rank correlation coefficients (ρ) are shown in each figure, all of which were not significant at the 5% level. Linear regression is shown in the figure, with solid lines indicating significance at the 5% level and dotted lines indicating non-significance.

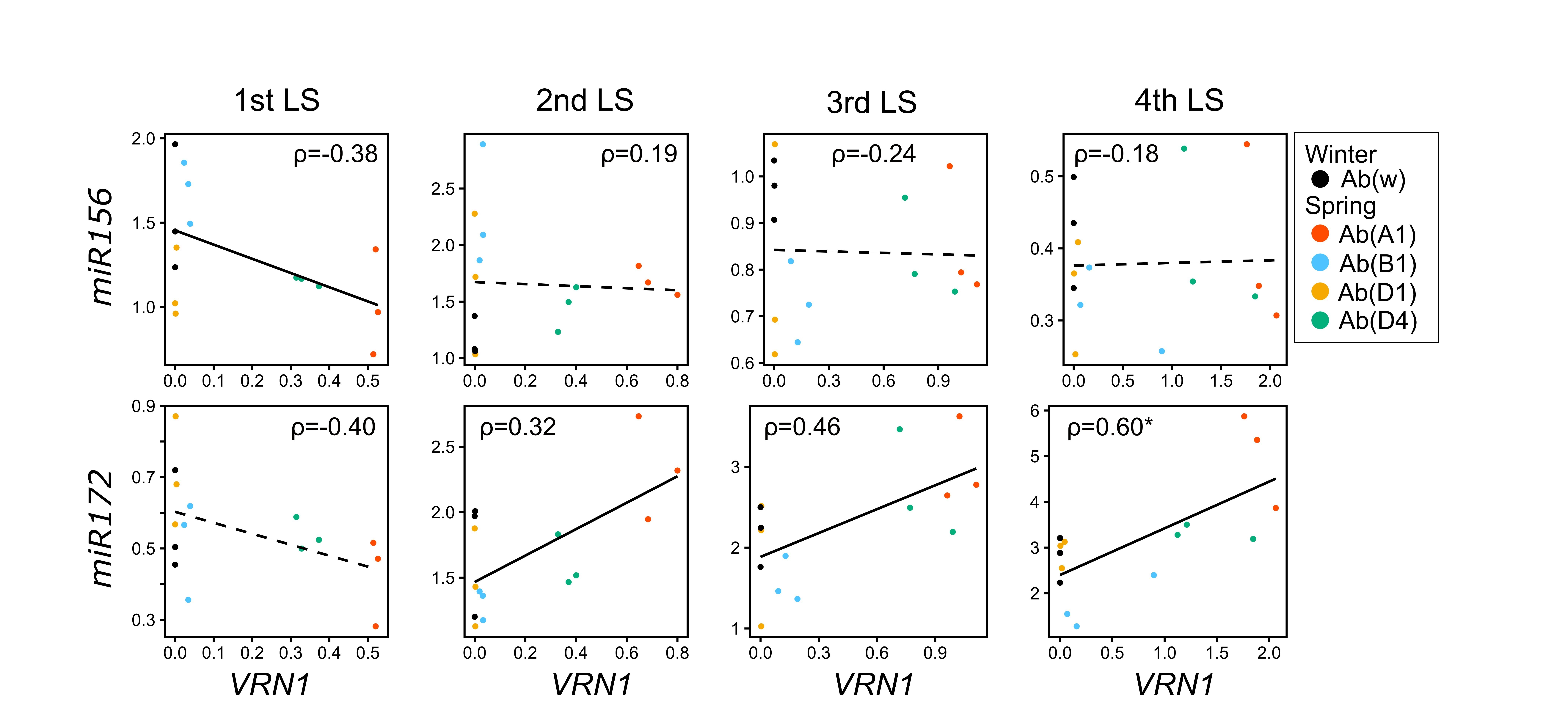

**Supplementary Fig. S6. Correlation of the expression levels of *VRN1* and miRNAs in the *VRN1* near-isogenic lines in each leaf stage.**

Each dot represents a biological replicate. Linear regression is shown in the figure, with solid lines indicating significance at the 5% level and dotted lines indicating non-significance. Spearman’s rank correlation coefficients (ρ) are shown in each figure with * (p < 0.05).

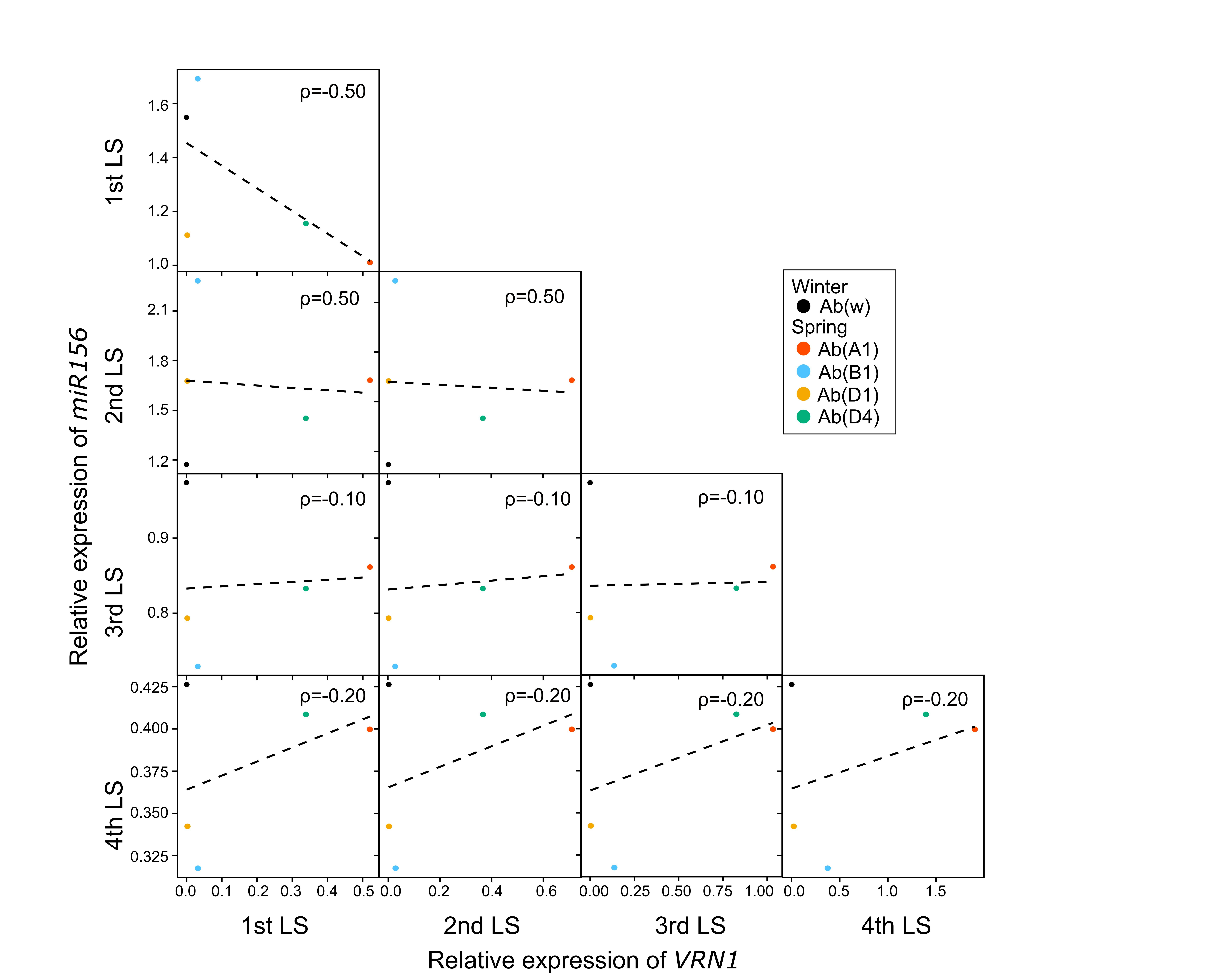

**Supplementary Fig. S7. Correlation between *VRN1* and *miR156* expression levels across leaf stages in the *VRN1* near-isogenic lines.**

Each dot represents the mean value in each variety. Linear regression is shown in the figure with dotted lines indicating non-significance at 5% level. Spearman’s rank correlation coefficients (ρ) are shown in each figure, all of which were not significant at the 5% level.

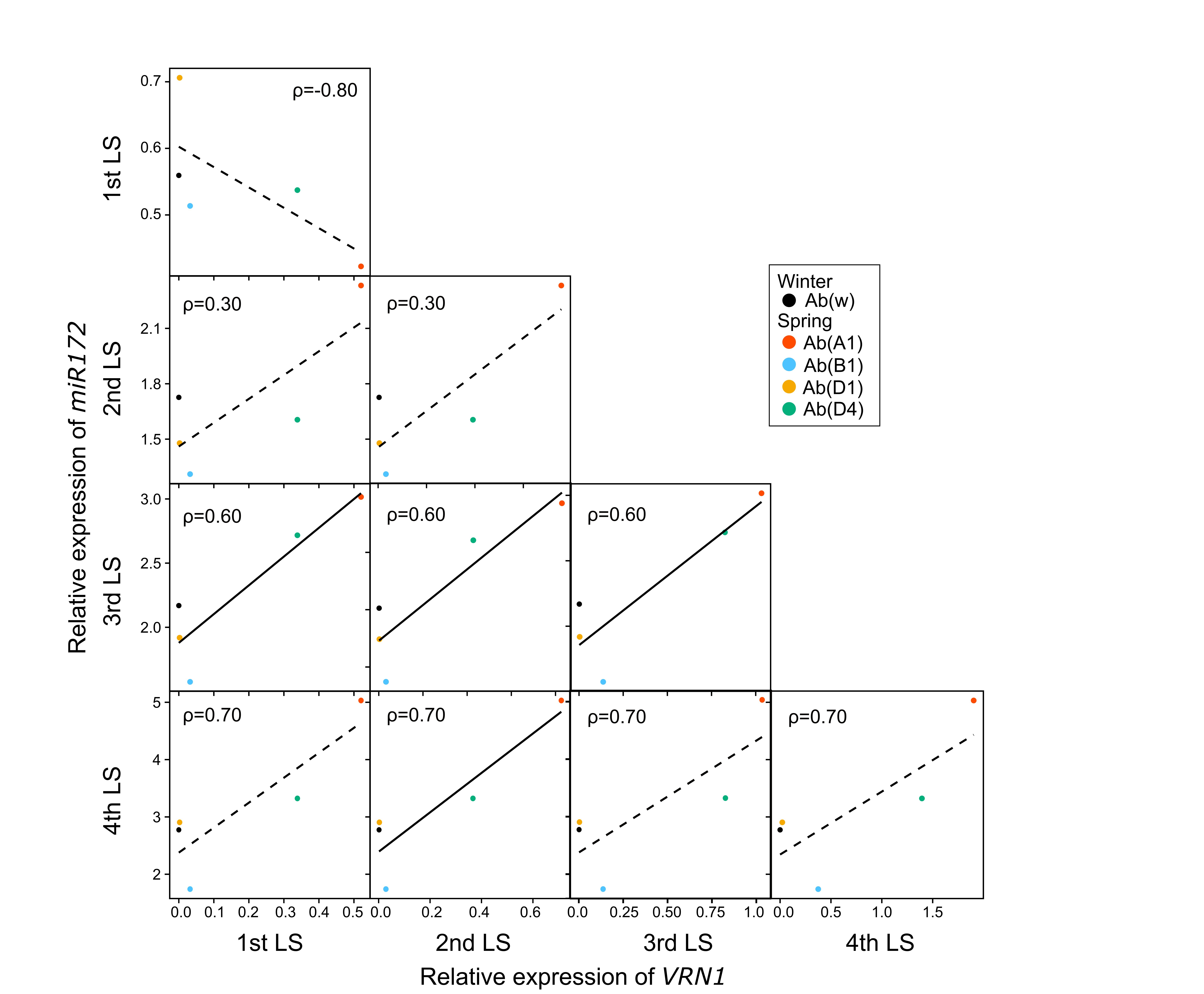

**Supplementary Fig. S8. Correlation between *VRN1* and *miR172* expression levels across leaf stage in the *VRN1* near-isogenic lines.**

Each dot represents the mean value in each variety. Linear regression is shown in the figure, with solid lines indicating significance at the 5% level and dotted lines indicating non-significance. Spearman’s rank correlation coefficients (ρ) are shown in each figure with * (p < 0.05), all of which were not significant at the 5% level.

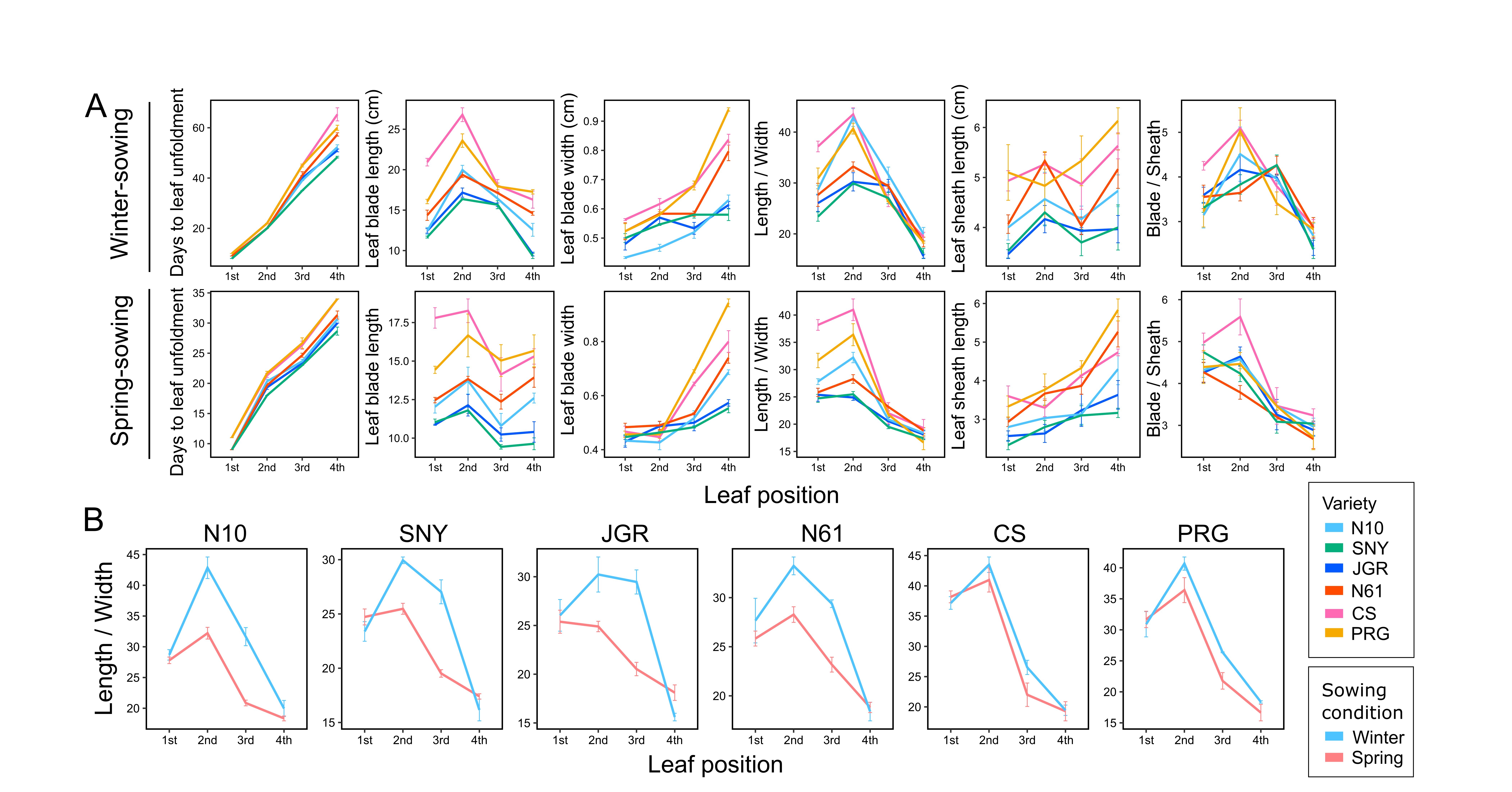

**Supplementary Fig. S9. Comparison of leaf morphological changes in the six common wheat varieties between the winter- and spring-sowing conditions in the field.**

(A) Measured values of the six traits, days to leaf stage, leaf blade length, leaf blade width, ratio of leaf blade length to width, leaf sheath length, and ratio of leaf blade length to leaf sheath blade, of the 1st to the 4th leaves under the winter-sowing (the upper) and the spring-sowing (the lower) conditions. Results are shown as the mean ± SE for each variety. n =3. (B) Comparison of the ratio of leaf blade length to width between the winter- and spring-sown conditions in each variety. Results are shown as the mean ± SE.

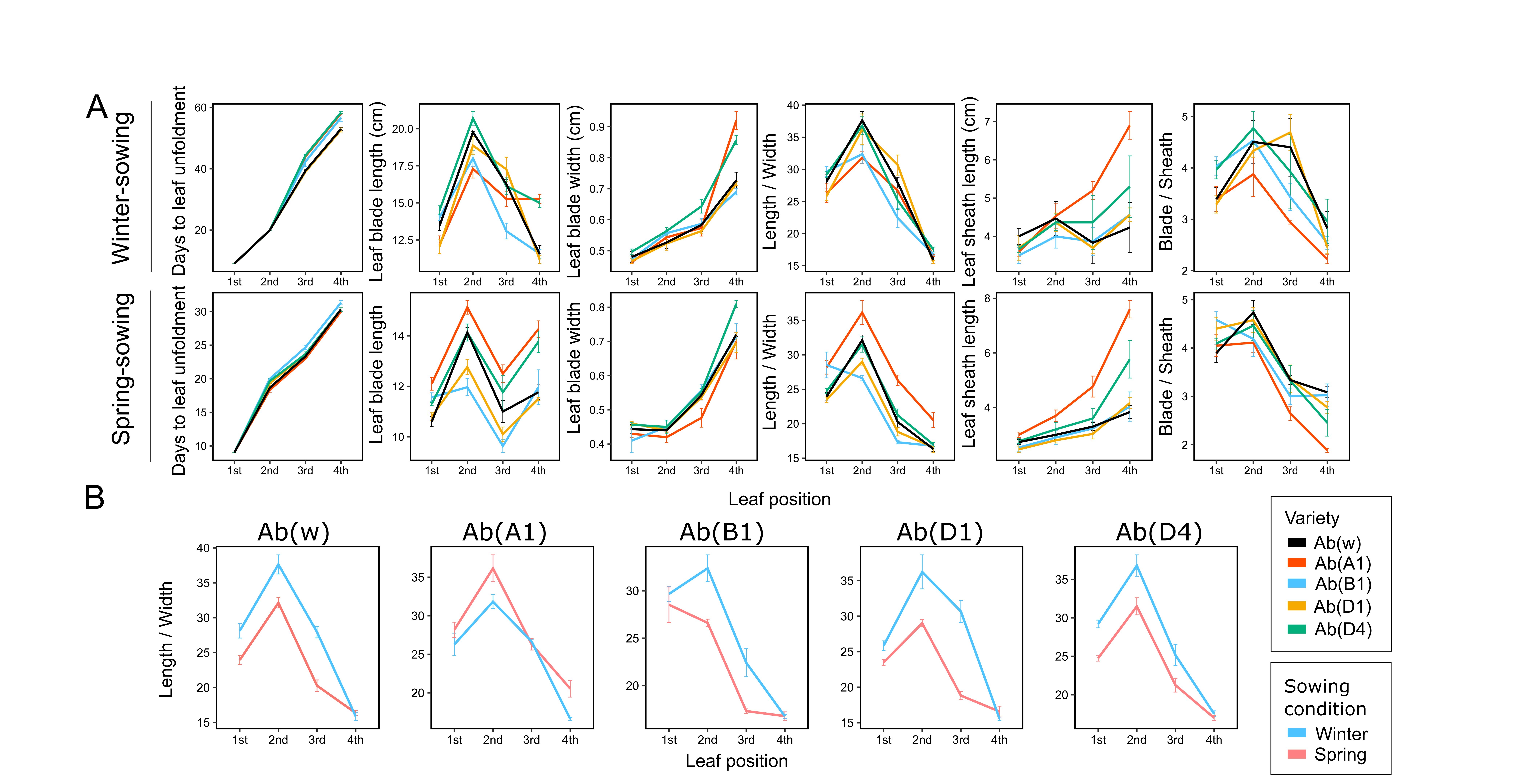

**Supplementary Fig. S10. Comparison of leaf morphological changes in the *VRN1* near-isogenic lines between in the winter- and spring-sown conditions in the field.**

(A) Measured values of the six traits, days to leaf stage, leaf blade length, leaf blade width, ratio of leaf blade length to width, leaf sheath length, and ratio of leaf blade length to leaf sheath blade, of the 1st to the 4th leaves under the winter-sowing (the upper) and the spring-sowing (the lower) conditions. Results are shown as the mean ± SE for each variety. n =3. (B) Comparison of the ratio of leaf blade length to width between the winter- and spring-sown conditions in each variety. Results are shown as the mean ± SE.

**Supplementary Table S1. Primers used in this study.**

| Name | Sequence | Length (bp) | Purpose | Reference |
| --- | --- | --- | --- | --- |
| WAP1-545L | GGAGAGGTCACTGCAGGAGGA | 21 | pPCR of VRN1 | Shimada et al., 2009 |
| WAP1-698R | GCCGCTGGATGAATGCTG | 18 |  |  |
| qPCR GAPDH F | AACTGCCTTGCTCCTCTTGC | 20 | qPCR ofGAPDH | This study |
| qPCR GAPDH R | ACCAGTGCTGCTTGGAATGA | 20 |  |  |
| SLO-miR156 | GTCTCCTCTGGTGCAGGGTCCGAGGTA  TTCGCACcagaggagACGTGCTC | 50 | RT^a^ of *miR156* | Debernardi et al., 2022 |
| SLO-miR172 | GTCTCCTCTGGTGCAGGGTCCGAGGTA  TTCGCACcagaggagACATGCAG | 50 | RT^a^ of *miR172* | Debernardi et al., 2022 |
| RT-miR156 | GGCGGTGACAGAAGAGAGT | 19 | qPCR for *miR156* | Debernardi et al., 2022 |
| RT-miR172 | GGCGGAGAATCTTGATGATG | 20 | qPCR for *miR172* | Debernardi et al., 2022 |
| Uni_MIRs | TGGTGCAGGGTCCGAGGTATT | 21 | qPCR for *miR156*  and *miR172* | Debernardi et al., 2022 |
| snoR101-Fw | GATGTCTTACACTTGATCTCTGAACTT | 27 | qPCR for *snoR101* | Debernardi et al., 2022 |
| snoR101-Rev | TGCATCAGGATTGATATAGTGTCC | 24 | RT^a^ of *snoR101* and qPCR  for *snoR101* | Debernardi et al., 2022 |

^a^ RT = reverse transcription
